# Structural basis of SNAPc-dependent snRNA transcription initiation by RNA polymerase II

**DOI:** 10.1101/2022.09.21.508861

**Authors:** Srinivasan Rengachari, Sandra Schilbach, Thangavelu Kaliyappan, Jerome Gouge, Kristina Zumer, Juliane Schwarz, Henning Urlaub, Christian Dienemann, Alessandro Vannini, Patrick Cramer

## Abstract

RNA Polymerase II (Pol II) carries out transcription of both protein-coding and non-coding genes. Whereas Pol II initiation at protein-coding genes has been studied in detail, Pol II initiation at non-coding genes such as small nuclear RNA (snRNA) genes is not understood at the structural level. Here we study Pol II initiation at snRNA gene promoters and show that the snRNA-activating protein complex (SNAPc) enables DNA opening and transcription initiation independent of TFIIE and TFIIH *in vitro*. We then resolve cryo-EM structures of the SNAPc-containing Pol II preinitiation complex (PIC) assembled on U1 and U5 snRNA promoters. The core of SNAPc binds two turns of DNA and recognizes the snRNA promoter-specific proximal sequence element (PSE) located upstream of the TATA box-binding protein TBP. Two extensions of SNAPc called wing-1 and wing-2 bind TFIIA and TFIIB, respectively, explaining how SNAPc directs Pol II to snRNA promoters. Comparison of structures of closed and open promoter complexes elucidates TFIIH-independent DNA opening. These results provide the structural basis of Pol II initiation at non-coding RNA gene promoters.

---

Transcription by RNA polymerase II (Pol II) has been structurally well studied for protein-coding genes that produce messenger RNA (mRNA)^1–4^. Pol II however also carries out transcription of non-coding small nuclear RNAs (snRNAs) that are an integral part of the pre-mRNA splicing machinery^5^. Pol II transcribes four of the five snRNAs, namely U1, U2, U4 and U5 snRNAs, whereas Pol III transcribes U6 snRNA^6^. In contrast to the Pol III-dependent snRNA promoter, Pol II-dependent snRNA promoters lack a TATA box motif^7^. To produce snRNAs, Pol II uses many of its accessory factors that are used for mRNA synthesis, but additionally requires specific factors for transcription initiation and elongation^8^.

Transcription initiation of snRNA genes relies on a specific factor called snRNA-activating protein complex (SNAPc). SNAPc binds a DNA motif in the upstream region of snRNA promoters, the so-called proximal sequence element, or PSE^9^. Human SNAPc contains five subunits – SNAPC1, SNAPC2, SNAPC3, SNAPC4 and SNAPC5. The subunits SNAPC1, SNAPC3 and SNAPC4 form the core of SNAPc^10^, of which SNAPC3 and SNAPC4 posess DNA-binding function^11,12^. The core subunits of SNAPc are conserved and have been characterized in Drosophila melanogaster and Trypanosoma brucei, where they are sufficient for activating snRNA transcription^13,14^. SNAPC2 and SNAPC5 however contribute to the stability and activity of SNAPc^10,15,16^.

The initiation of SNAPc-regulated Pol II snRNA transcription was reported to rely on the general transcription factors (GTFs) TBP, TFIIA, TFIIB, TFIIE and TFIIF^17,18^. The role of TFIIH in Pol II snRNA transcription remains unclear^17^, although TFIIH is known to be required for DNA opening at promoters of protein-coding genes^19^. The structure of SNAPc and its molecular interactions with the Pol II pre-initiation complex (PIC) are also unknown. As a consequence, the structural basis and the mechanism of snRNA transcription initiation remains to be uncovered. Here we report structures of SNAPc-containing Pol II PICs bound to U1 and U5 snRNA promoters. Our results show how SNAPc is structured, how it recognizes the PSE, and how it positions a core Pol II PIC on snRNA promoters for DNA opening and transcription initiation. More generally, this work adds to our understanding of the evolution of the three eukaryotic transcription systems.

## RESULTS

### Preparation of functional SNAPc

We prepared two variants of recombinant human SNAPc, namely SNAPc-FL containing all full-length subunits and SNAPc-core^10^, containing SNAPC1, SNAPC3, SNAPC4 (residues 1-516) and SNAPC5 (**Figure 1a**) (Methods). Both purified SNAPc variants were able to bind U1 and U5 snRNA promoter DNA (RNU1 and RNU5), both in the absence and in the presence of TBP and TFIIB in an electrophoretic mobility shift assay (EMSA) (**Figure 1b**). EMSA also showed that the SNAPc variants could facilitate binding of TBP to snRNA promoters that lack a TATA box (**Figure 1b, 2a**), consistent with previous studies^20^.

**Figure 1.**
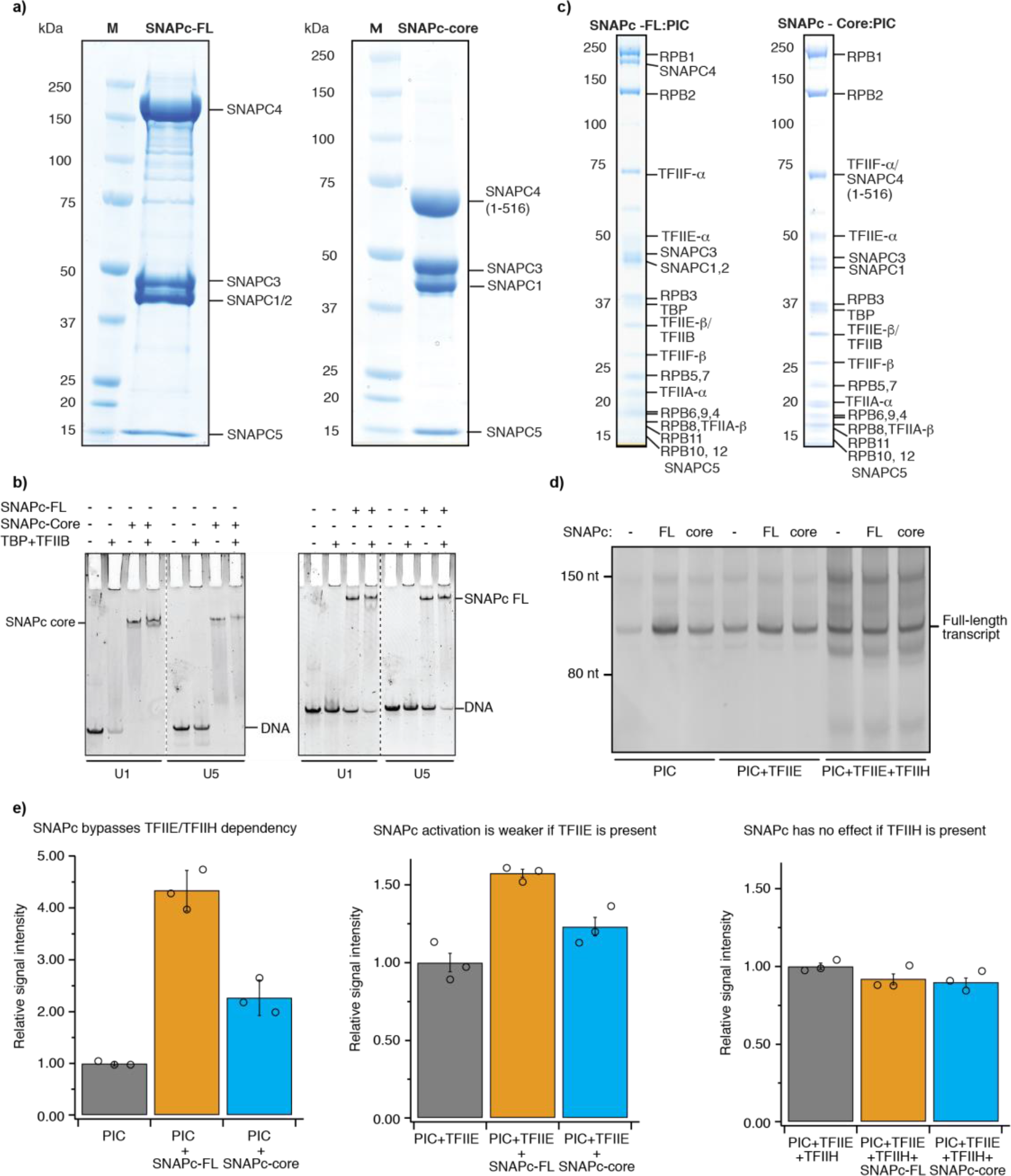
Preparation of SNAPc-containing Pol II PIC on non-coding RNA promoters. a) SDS-PAGE analysis of SNAPc variants (FL, core) purified to homogeneity. b) EMSA shows the binding of SNAPc (± TBP, TFIIB) to U1 and U5 promoter DNA. The presence of SNAPc stabilises the binding of TBP-TFIIB to snRNA promoters. c) SDS-PAGE analysis of SNAPc containing Pol II PIC variants isolated through a sucrose gradient ultracentrifugation. d) In vitro transcription assay showing the relative influence of SNAPc variants on Pol II snRNA transcription with different combinations of GTFs’. e) Histogram plots representing the quantification (Methods) of full-length transcripts from the *in vitro* transcription assay in panel d.

**Figure 2.**
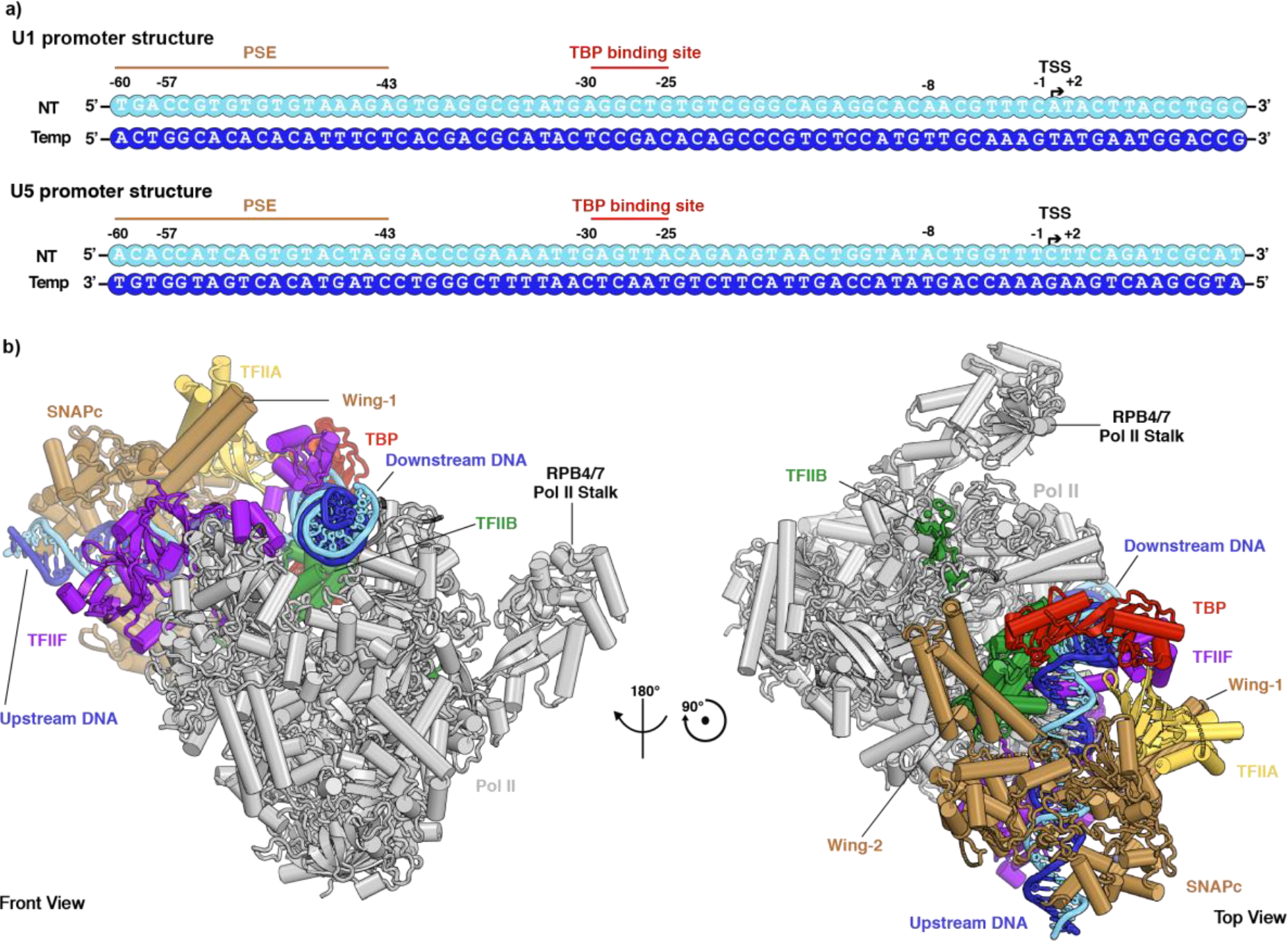
Overall structure of SNAPc-containing Pol II PIC. a) Schematic 2D representation of the U1 and U5 promoter sequences highlighting the binding motifs of the initiation machinery as observed in the cryo-EM structure: PSE (SNAPc), TBP binding site (TBP) and TSS (Pol II). The transcription start site (TSS) is denoted +1 and negative and positive numbers indicate upstream and downstream positions. b) Cartoon representation of the SNAPc-containing Pol II PIC as viewed from the front and top. The colour codes for Pol II and the GTFs’ are consistently used throughout.

To test whether recombinant SNAPc could mediate Pol II transcription initiation from snRNA gene promoters, we used an *in vitro* transcription assay. The assay showed that Pol II could initiate transcription from a U1 promoter in the presence of TBP, TFIIA, TFIIB and TFIIF and was stimulated ~4.5 fold and ~2.5 fold by the addition of SNAPc-FL or SNAPc-core, respectively (**Figure 1d, e**). Addition of TFIIE reduced this increase in transcription activity to ~1.5 fold and ~1.2 fold for SNAPc-FL and SNAPc-core, respectively (**Figure 1e**), suggesting that TFIIE is not required for SNAPc-dependent transcription initiation and rather inhibitory in our biochemical system. Further addition of TFIIH did override the stimulatory effect of SNAPc and led to formation of non-specific transcripts at multiple sites (**Figure 1d**). In conclusion, our recombinant SNAPc variants stimulate Pol II transcription initiation from snRNA gene promoters in the absence of TFIIE and TFIIH *in vitro*.

## Cryo-EM analysis of SNAPc-containing PICs

Based on these observations we reconstituted a functional SNAPc-containing Pol II PIC on a U1 promoter DNA (Methods). We incubated SNAPc-core and S. scrofa Pol II (99.9% identical to human Pol II) with human TBP, TFIIA, TFIIB, TFIIE and TFIIF, and subjected the resulting complex to sucrose-gradient ultracentrifugation. Peak fractions contained apparent stochiometric amounts of Pol II, SNAPc-core and the general transcription factors, indicating formation of a stable 24-subunit SNAPc-containing PIC (**Figure 1c**). The sample was crosslinked^21^ and subjected to cryo-EM analysis (Methods). Initial trials showed that the PIC containing the SNAPc-FL variant was less stable (**Figure 1c**), whereas the PIC containing SNAPc-core was stable and suited for cryo-EM analysis, leading to a high-resolution single particle dataset (**Extended Data Table 1**).

Reconstructions from 3D classification of this dataset showed two distinct particle classes of the SNAPc-containing Pol II PIC (**Extended Data Figure 1**). Further 3D classification and refinement identified these two states as the closed promoter complex (CC) and the open promoter complex (OC) states of the PIC. We obtained structures of the CC and OC states at an overall resolution of 3.4 Å and 3.0 Å, respectively (**Extended Data Figure 1, 3**). None of our maps revealed density for TFIIE, consistent with our in vitro transcription assays that showed TFIIE was not required for initiation (**Figure 1d, e**). Densities for SNAPc and upstream DNA containing the PSE were improved by focussed 3D classification and masked refinements. The local resolution for this region was 3.5 Å for the OC state (**Extended Data Figure 1, 3)**.

In an effort to obtain a high-resolution structure of SNAPc, we additionally reconstituted a SNAPc-containing Pol II PIC on a DNA that was based on the U5 promoter sequence (Methods). The resulting cryo-EM dataset enabled refinement of the SNAPc-containing PIC in the CC state at an overall resolution of 3.0 Å and with the local map of the upstream region extending to 3.2 Å (**Extended Data Figure 2, 3**). The local map enabled building of an atomic model for SNAPc and PSE-containing upstream DNA (Methods). We then combined the resulting model with the known high-resolution structures of mammalian Pol II PIC in CC and OC states^22^. After manual adjustment, refined structures of the SNAPc-containing PIC in the CC and OC states containing the U1 and U5 promoters showed good stereochemistry resulting in a total of three structures (**Extended Data Table 1**).

### Overall structure of SNAPc-containing PIC

The overall structure of the SNAPc-containing Pol II PIC shows that SNAPc binds the promoter DNA upstream of TBP (**Figure 2**). SNAPc recognizes the PSE motif and interacts with TFIIA and TFIIB. Despite these multiple interactions, the presence of SNAPc does not alter the canonical core PIC structure in any substantial way^22^. TBP binds to the AGGCTG sequence at register –30 to –25 bp (**Figure 2a**) of the TATA-less U1 promoter and bends the DNA by 90° similar to what is observed in a TBP-TATA DNA complex^23,24^ (**Extended Data Figure 4a**). In the following, we will first describe the SNAPc structure and SNAPc-DNA interactions based on the U5-containing CC structure that is resolved at the highest resolution. We will then describe promoter opening based on the CC and OC structures of the U1-containing PIC.

### SNAPc structure contains two protruding wings

The high-resolution structure of the SNAPc core bound to the U5 promoter shows how the subunits SNAPC1, SNAPC3 and SNAPC4 fold and interact (**Figure 3**). SNAPC1 possesses an N-terminal VHS/ENTH-like domain^25^ that forms a mainly helical structure (**Extended Data Figure 4b**). SNAPC3 is saddle-shaped with a central ‘ubiquitin-like domain’ (ULD) and additional α-helices and β-strands (**Extended Data Figure 4c**). Consistent with biochemical studies^26^, SNAPC3 contains two zinc fingers (ZF-1 and ZF-2). ZF-1 is a C2H2 type zinc finger with residues Cys221, His313, Cys317 and His319 coordinating a Zn^2+^ ion (**Extended Data Figure 5f**). ZF-2 is a C4 type zinc finger with residues Cys354, Cys357, Cys380, Cys383 coordinating another Zn^2+^ ion (**Extended Data Figure 5g**). SNAPC4 contains four complete repeats (R1-R4) and a half repeat (Rh) of the Myb domain^12^, of which we observe Rh, R1 and R2 (residues 274-398) (**Figure 3b**, **Extended Data Figure 4d**). R1 and R2 contain three helices forming canonical helix-turn-helix folds. The SNAPc core is stabilized by intricate interactions of SNAPC3 with both SNAPC1 and SNAPC4. The N-terminal region of SNAPC3 interacts mainly with SNAPC1, burying a surface area of ~1640 Å^2^. The C-terminal region of SNAPC3 binds SNAPC4 and buries ~3010 Å^2^. A total of four subunit interfaces are formed based on hydrophobic interactions, salt bridges and polar contacts (**Figure 3c-f, Extended Data Figure 7**).

**Figure 3.**
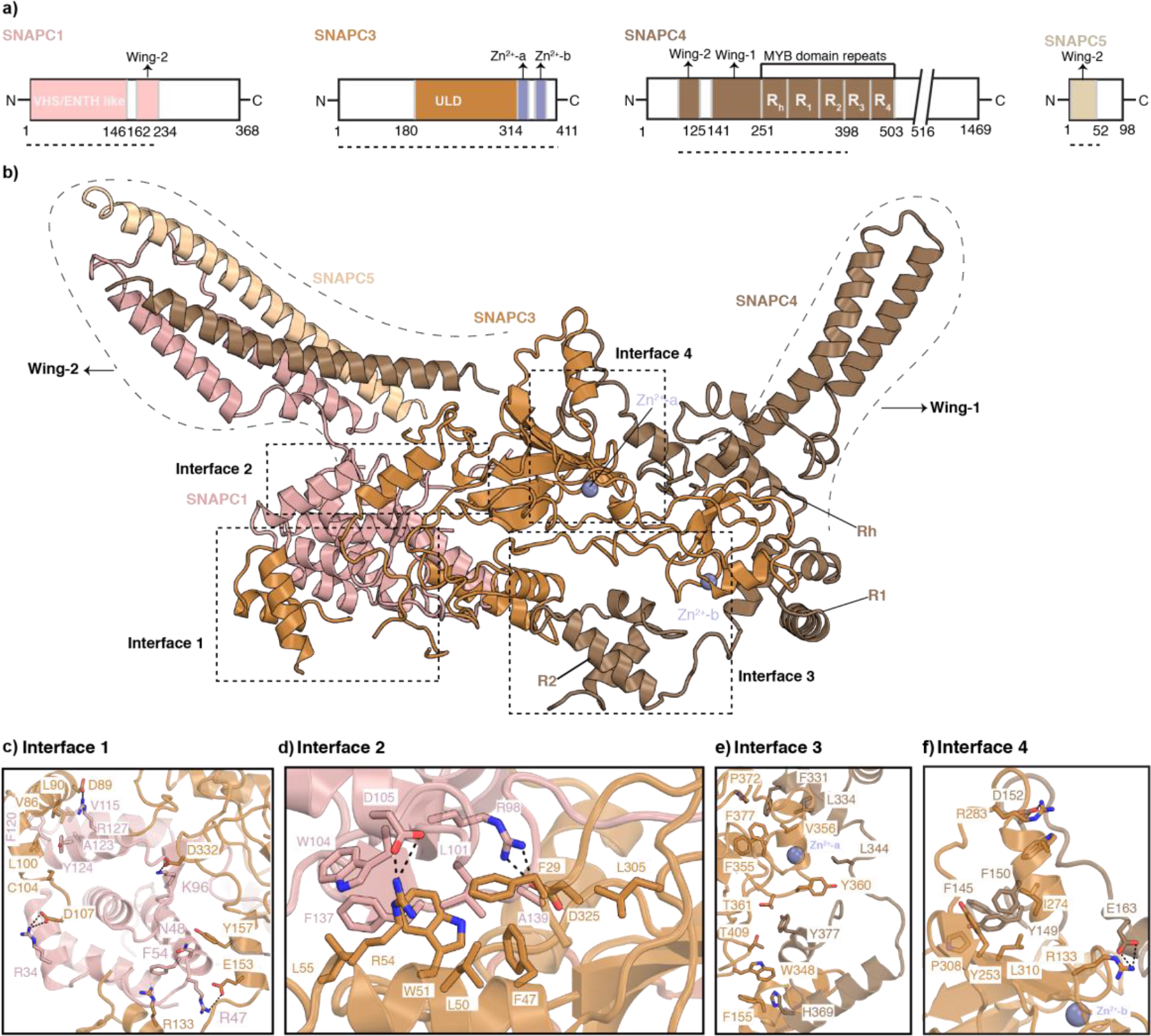
Structure of SNAPc. a) 2D-domain schematics of individual SNAPc subunits. The regions visible in the 3D structure are marked by dotted-lines. b) SNAPc structure in cartoon representation. Domain nomenclature and colours are used as described in panel a. Dashed boxes indicate the interfaces between the subunits. c, d) Close up view of interfaces 1 and 2 that are formed between SNAPC1 (pink) and SNAPC3 (orange). The residues V115, F120, A123, Y124 of SNAPC1 and V86, L90, L100, C104 of SNAPC3 form mainly hydrophobic interactions, whereas ionic interactions are formed between R34, R47, K96, R128 of SNAPC1 and D89, D107, E153, D332 of SNAPC3. F54 of SNAPC1 and R133 of SNAPC3 form a cation-pi interaction and N49 of SNAPC1 and Y157 of SNAPC3 form polar contacts. Similarly in interface 2: SNAPC1 L101, W104, F137 and A139 form hydrophobic contacts with F47, L50, W51, L55 and L305 of SNAPC3. Salt-bridges involving R98, D105 of SNAPC1 and R54 and D325 of SNAPC3 fortify interface 2. e, f) Interfaces 3 and 4 between SNAPC3 (orange) and SNAPC4 (chestnut brown). In interface 3, SNAPC3 residues F155, W348, F355, V356, Y360, T361, P372, F377, T409 form the bulk of hydrophobic contacts with F331, L334, L344 and H369 of SNAPC4 (Figure 3e). Likewise in interface 4 the residues Y253, I274, W277, P308 and L310 make hydrophobic contacts with the amino acids F140, Y149, F150, F176 of SNAPC4. Additional salt bridges are formed by R133, R283 of SNAPC3 with D152 and E153 of SNAPC4. The Zn-fingers (ZF-1, ZF-2) of SNAPC3 are in close proximity to the interfaces 3 and 4, and would be important for the structural integrity of this complex. The residues involved in these protein-protein interaction surfaces are highly conserved across metazoans (Extended Data Figure 7).

SNAPc also contains two protrusions that we refer to as ‘wing-1’ and ‘wing-2’. The wing-1 of SNAPc consists of a pair of helices that precede the Rh region of SNAPC4 (residues 184-256). The wing-2 of SNAPc is a four-helix bundle that is formed by two helices of SNAPC1 (residues 162-234) and one helix each of SNAPC4 (residues 81-125) and SNAPC5 (residues 1-51) (**Extended Data Figure 4e**). Although the resolution in wing-2 is limited due to mobility, AlphaFold2 prediction^27^ and prior biochemical studies^16^ led to a reliable model of wing-2 that we confirmed by crosslinking mass-spectrometry (**Extended Data Figure 5k, 6**). In conclusion, these efforts provided the structure of SNAPc, which contains a three-subunit core and two protruding wings extending from the core.

### SNAPc core recognizes the snRNA promoter

Our U5-containing CC structure also reveals details of how SNAPc binds the PSE motif in promoter DNA (**Figure 4a**). The SNAPc core binds to the PSE motif through its subunits SNAPC3 and SNAPC4 (**Extended Data Figure 8a**), consistent with biochemical data^11,28^. SNAPc contacts promoter DNA 8 bp upstream of the proximal edge of the TBP-binding site (**Figure 4b, c**). The register of the modelled snRNA promoter is defined by the nucleotide on the non-template (NT) strand at the upstream edge of TBP binding site starting at –30, ascending in the 5’ to 3’ direction. SNAPC3 and SNAPC4 both bind this region through contacts with the DNA backbone and bases on both strands of the PSE (**Extended Data Figure 8a**). DNA binding occurs both to the major and minor grooves. SNAPC3 inserts its helix α8 into the major groove and forms multiple contacts with DNA. K199 forms salt bridges with the backbone phosphates of nucleotide G9 on the template strand. The residue K194 of the same helix forms ionic interactions with the O6 atom of the nucleotide bases G –42 and G –43 of the NT strand. Further downstream, H198 establishes hydrophobic contacts with the nucleotide base T –45 on the template strand (**Figure 4b**).

**Figure 4.**
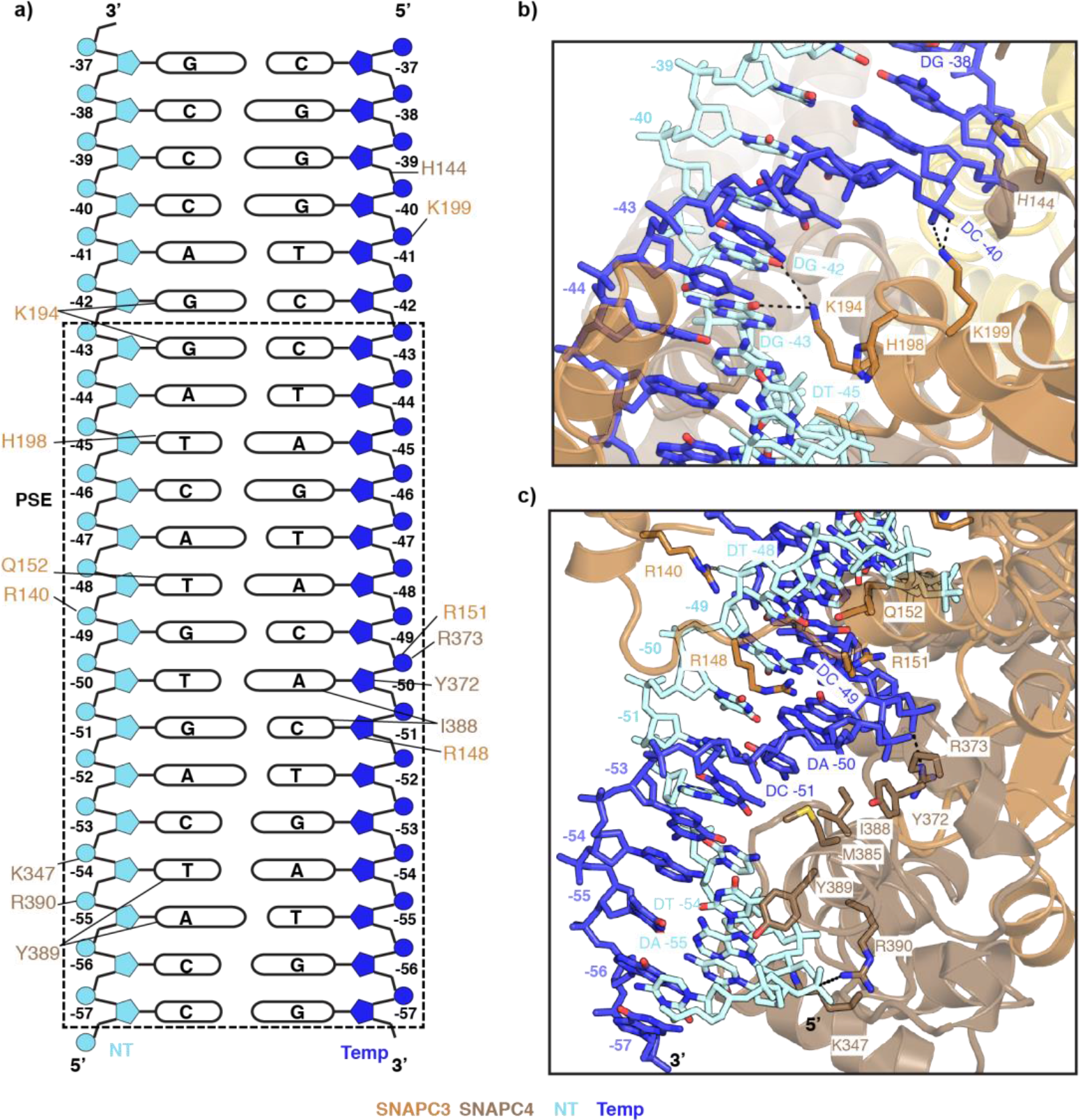
SNAPc-DNA interactions. a) Schematic view of the protein-DNA interactions between SNAPc and the PSE motif. Residues interacting with specific regions of the DNA as described in the text are indicated by lines. In panels b and c, nucleotide residues are numbered in atomic colour to indicate the strand and the DNA register b) DNA-protein interaction network on the preceding major and minor grooves (register: −46- to −35) of PSE as bound by SNAPc subunits SNAPC3 and SNAPC4. Colour codes are used uniformly in all panels. c) Close up view of the first major and minor groove (register: −57 to −47) interactions between SNAPc and the PSE motif on U5 promoter. The SNAPc subunits are represented as cartoon, whereas the interacting amino acid sidechain residues, DNA chains are depicted as sticks with atomic colours. Dashed lines indicate ionic interactions.

Since most of these protein-DNA contacts are to the DNA backbone, the question arises how SNAPc can recognize the PSE. Our structure suggests that recognition is at least partially achieved by indirect readout. In particular, the DNA major groove is locally distorted at the PSE and differs from canonical B-DNA at registers –51 to –41 (**Extended Data Figure 8b**). At the position where SNAP3 helix α8 is inserted into the major groove, the duplex geometry resembles A-form DNA^29^ (**Extended Data Figure 8c**). This deviation is also reflected in the minor grooves upstream and downstream of this site (**Extended Data Figure 8a, d**).

SNAPc also binds the minor groove of DNA with subunits SNAPC3 and SNAPC4. Q152 of SNAPC3 a forms hydrogen bond with the nucleotide base T –48 of NT strand while SNAPC4 residue Y372 interacts hydrophobically with the C3 atom of backbone sugar of the nucleotide base A –50 of the template strand. Arginine residues R148 and R151 of SNAPC3 and R373 of SNAPC4 form salt bridges with the DNA backbone (**Figure 4c**). Our structure also shows that the SNAPC4 Myb repeat R2 binds DNA via its helix α15 that contacts the anterior major groove, and early biochemical studies indicated that the Myb repeats R3 and R4 are involved in DNA binding^12^. I388 establishes hydrophobic interactions with the nucleotide base A –50 and the C5 atom of nucleotide C –51 on the template strand. The neighbouring Y389 residue forms hydrogen bonds with the N7 atom of A –55 and hydrophobic interaction with T −54 of the NT strand (**Figure 4c**). The residues K347, R373 and R390 of SNAPC4 interact with the DNA backbone. Although biochemical studies had identified SNAPC3 and SNAPC4 as poor DNA binders when investigated in isolation^10,11^, our results suggest that formation of the SNAPc complex with its intricate interactions between these two subunits enables tight binding of the PSE which explains how SNAPc recognizes the snRNA promoter.

### The wings of SNAPc bind TFIIA and TFIIB

SNAPc also interacts with TFIIA and TFIIB that flank TBP in the PIC (**Figures 5, Extended Data Figure 8a**). Whereas wing-1 of SNAPc binds to TFIIA, wing-2 binds TFIIB (**Extended Data Figure 8a**). SNAPc interaction with TFIIA and TFIIB involves three interfaces that we call A, B and C. In interface A, the wing-1 of SNAPC4 (helices α4, α5) slides under the four-helix bundle of TFIIA like a wedge, stabilizing the flexible bundle region (**Figure 5a**). SNAPC4 additionally interacts with the β-barrel of TFIIA to form interface B (**Figure 5a**). Interfaces A and B are formed by a combination of hydrophobic interactions, salt bridges and polar contacts. Incidentally, the TFIIA bundle has also been shown to interact with TAF4 and TAF12 in lobe B of the multisubunit TFIID complex that, like SNAPc, is important for promoter recognition^30^. Interface C is formed between wing-2 and the C-terminal cyclin fold of the TFIIB core (**Figure 5b**). The wing-2 helices from SNAPC1 and SNAPC5 form contacts with the terminal α-helix of the TFIIB core. Interface C stabilizes the TFIIB core, which was suggested to play a key role in the activation of snRNA transcription initiation^7^. Together, SNAPc wing-1 and wing-2 bind TFIIA and TFIIB, respectively, to position the core PIC with respect to SNAPc and the PSE promoter element.

**Figure 5.**
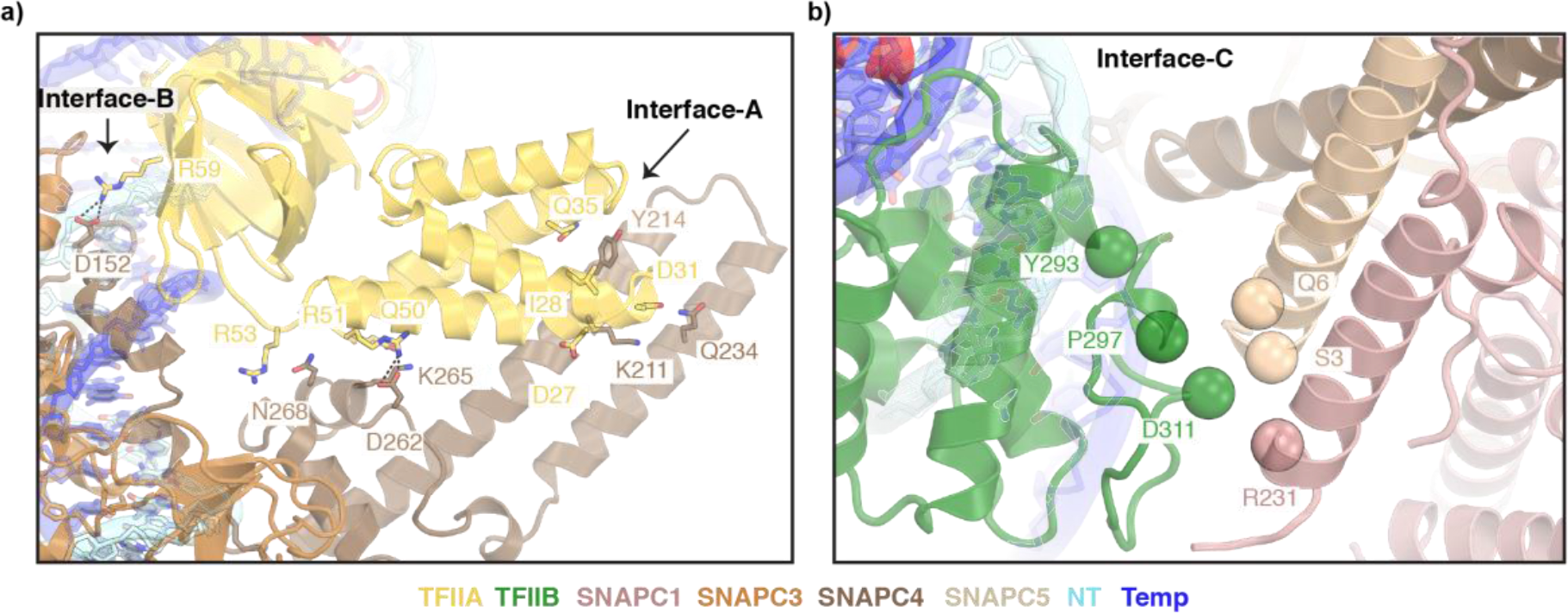
SNAPc-general transcription factors interaction. a) Close up view of wing-1:TFIIA interaction. The amino acid residues involved in the formation of interfaces A and B between TFIIA (yellow orange) and SNAPC4 (chestnut brown) are represented as sticks. Dashed lines indicate salt-bridges. b) Zoomed in view of the interface C formed between wing-2 and TFIIB C-terminal cyclin fold. The Cα atoms of putative residues forming the interaction surface are represented as spheres.

### Promoter DNA opening

Comparison of our CC and OC structures bound to the U1 promoter provides insights into the mechanism of TFIIE- and TFIIH-independent DNA opening (**Figure 6**). Overall, closed and open U1 promoter DNA follow trajectories within the Pol II cleft that are comparable to those observed for protein-coding promoter DNA in PIC structure^22^. Also, as observed in PIC structures lacking SNAPc^2,22^, the OC state is associated with a closed Pol II clamp and an ordered B-reader and B-linker elements in TFIIB (**Figure 6b**). However, DNA opening can also be achieved spontaneously in the absence of TFIIE and TFIIH at some protein-coding genes in yeast^31^, and such spontaneous opening depends on the DNA duplex stability around the transcription start site (TSS)^32^. Studies in yeast Pol II have further shown that an AT-rich sequence increases the propensity of spontaneous promoter opening during transcription intiation^31^. Similarly, we find that promoter sequences of snRNA-encoding genes are AT-rich in the initially melted region (IMR) spanning positions –8 to +2 around the TSS (position +1) (**Extended Data Figure 8e**). We propose that the AT-rich nature of the IMR enables spontaneous DNA opening of the U1 promoter upon PIC binding. In summary, these results suggest that DNA opening of snRNA gene promoters and the spontaneously melted protein-coding genes rely on easily melting regions around the TSS and use similar mechanisms.

**Figure 6.**
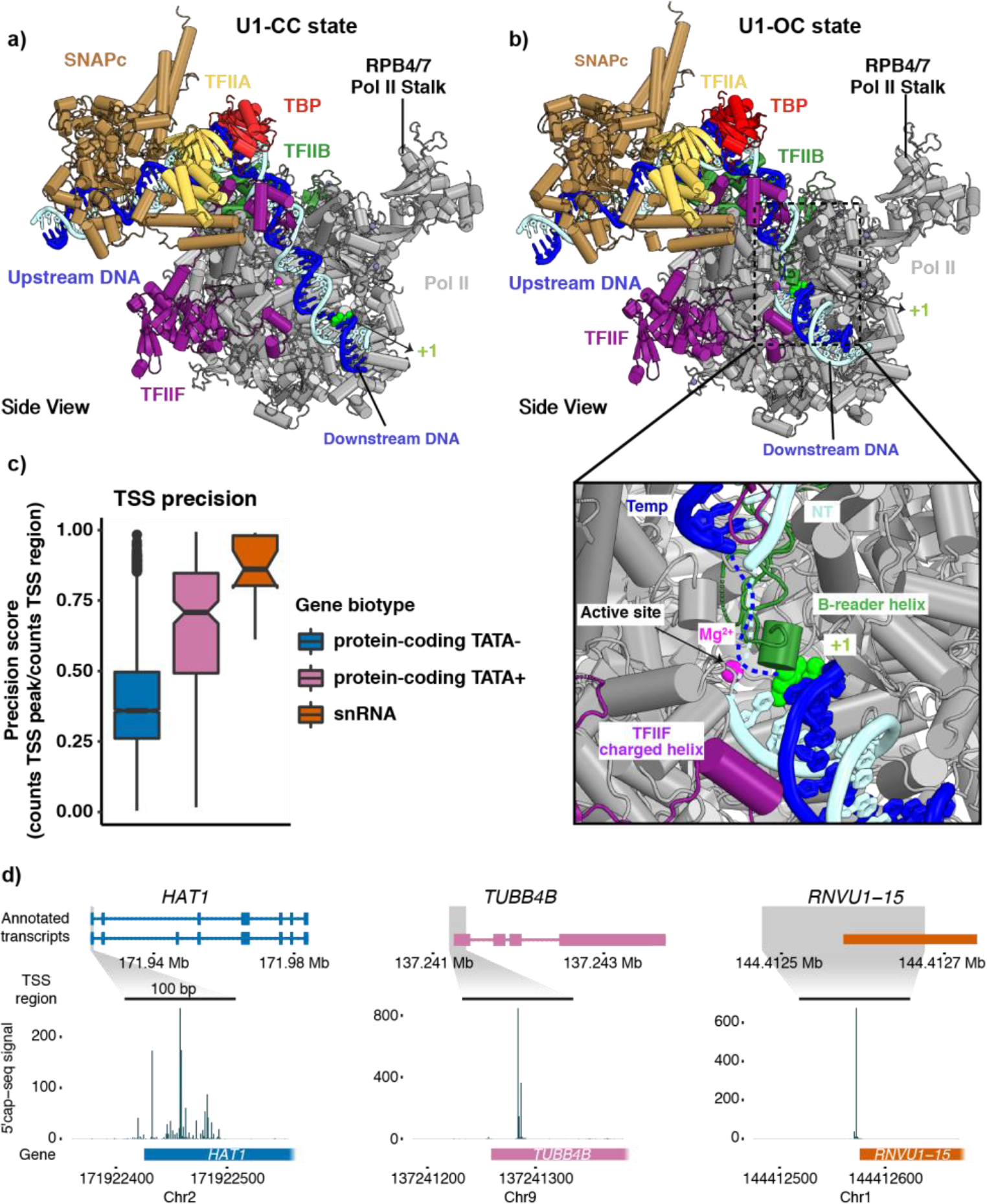
Promoter opening. a) Structure of SNAPc-containing Pol II PIC bound to U1 promoter in closed promoter complex (CC) state. The subunits are coloured as in Figure 1. The nucleotide residue at the TSS (+1) on the template strand (blue) is represented as spheres (green). The Pol II active site metal ion A is depicted as a magenta sphere. b) Structure of SNAPc-containing Pol II PIC bound to U1 promoter in open promoter complex (OC) state. The inset represents a zoom into the active center containing open promoter DNA. The catalytic Mg^2+^ ion at the active site is represented as a magenta sphere. The B-reader helix of TFIIB and the charged helix of TFIIF are highlighted alongside the +1 nucleotide residue represented as a sphere (green). c) Box plots showing TSS precision of protein-coding and snRNA genes (N=18) transcribed by Pol II in cells. Protein-coding genes are sub-grouped based on promoter sequence into TATA-less (TATA–, N=4521) and TATA-containing (TATA+, N=200) subsets. The thickened line represents the median value, the hinges correspond to the first and third quartiles, and the notches extend to 1.58 times the inter-quartile range divided by the square root of N. The whiskers represent the largest or smallest value within the 1.5 times inter-quartile range from the hinge, outliers are shown in black. The precision scores were determined from published 5’cap-seq data^35^ (Methods). d) Annotated transcripts of representative examples from subsets in 6C and genome browser views showing 5’cap-seq signal in the magnified region (± 100 bp) centered at the main TSS peak. The annotated gene region is show below the views and only sense strand signal is shown.

### Definition of the transcription start site

We observe 19 nucleotides of the DNA template strand spanning from the TBP-binding site to the upstream edge of the DNA bubble (at position −12). The templating nucleotide in open promoter DNA reaches the active site of Pol II ~30 nucleotides downstream of the upstream edge of the TBP-binding site (**Figures 6a, b**). The DNA strands forming the open DNA bubble are mobile, leading to a weakly resolved map. Subsequently, 12 nucleotides further downstream, we observe T +1 of the template strand immediately downstream of the catalytic Mg^2+^ ion at the active site. This posits residue G –1 as the template for RNA synthesis. The CA dinucleotide is the signature of the Initiator sequence (Inr)^33^ and is located at register –1 and +1 of the non-template strand. This observation suggests that the TSS position is defined by a fixed distance from the site of TBP binding, as is known for protein-coding human genes that have their TSS within a window of 28-33 bp downstream of the TATA box^34^. Since we also observe a fixed position of SNAPc with respect to TBP, the TSS is apparently set by a fixed distance from the PSE in snRNA promoters.

These observations suggest that Pol II transcription would initiate from a TSS that is rather precise *in vivo*. To investigate this, we identified the main TSSs and determined their ‘TSS precision scores’ from a reanalysis of 5’-capped RNA sequencing data^35^ for both mRNA and snRNA encoding genes with a constitutive first or a single exon (Methods). A maximum precision score of 1 means that all transcripts initiate at the main TSS (±2 bp). Indeed we find that Pol II snRNA transcription generally initiates in this narrow, 5-bp window with a high median precision score of 0.86, as exemplified by the *RNVU1-15* promoter (**Figure 6c**). In contrast, Pol II initiates transcription less precisely at TATA-less mRNA promoters, as shown by a median precision score of 0.36, as exemplified by the *HAT1* promoter. Pol II also initiates mRNA transcription more precisely when promoter DNA contains a TATA box motif, with a median precision score of 0.71, as exemplified by the *TUBB4B* promoter (**Figure 6c**). These large differences in TSS precision are also observed in genome browser views of representative promoters (**Figure 6d**). The observed high TSS precision of snRNA promoters is consistent with our model that SNAPc defines TSS position. In summary, SNAPc binding to the PSE likely serves as a ruler for positioning of TBP at TATA-less snRNA promoters, leading to initiation at a defined distance downstream of the PSE.

## DISCUSSION

Here we report structures of SNAPc-containing Pol II PICs on two different snRNA gene promoters and in two different states, the CC and OC states. Together with biochemical results and published literature, our structures suggest the mechanism of SNAPc-mediated snRNA transcription initiation by Pol II (**Figure 7**). SNAPc uses its conserved core to recognize the PSE motif in snRNA promoters, whereas its two wings position TFIIA and TFIIB. Since TFIIA and TFIIB form a rigid complex with TBP, SNAPc can indirectly position TBP at a defined location on snRNA promoters despite the absence of a consensus TATA box motif. This is consistent with the evidence that TFIIB-TBP complexes can be effectively recruited to snRNA promoters exclusively as part of a ternary TFIIA-TFIIB-TBP complex^18^. Positioning of the TFIIA-TFIIB-TBP complex on promoter DNA in turn recruits the Pol II-TFIIF complex to the IMR of the promoter. The low DNA duplex stability at the IMR enables spontaneous DNA opening and occurs with the use of binding energy independent of TFIIE and TFIIH. The emerging DNA template strand then binds in the Pol II active center cleft and RNA chain synthesis is initiated at an Inr dinucleotide CA^33^, thereby setting the TSS at a defined distance from the PSE.

**Figure 7.**
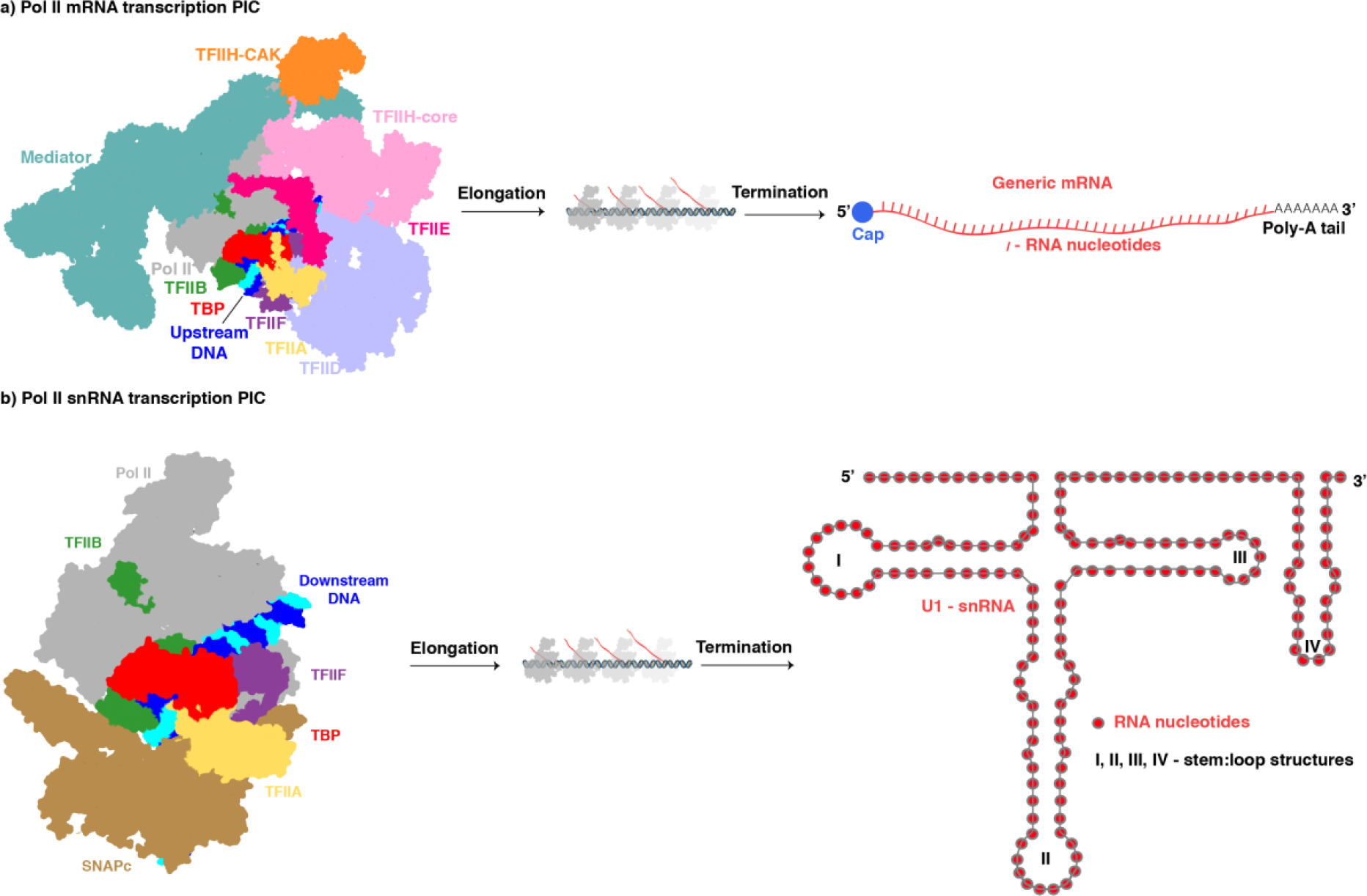
Comparisons of Pol II PICs for mRNA and snRNA synthesis. a) The Pol II PIC on protein coding genes bound to its elaborate array of initiation factors such as TFIIA, TFIIB, TFIID, TFIIE, TFIIF, TFIIH and Mediator complex. b) The Pol II PIC for snRNA transcription requires SNAPc but not TFIIE, TFIIH and Mediator to initiate transcription.

Comparison of our results with published data also provides insights into the evolution of the three different eukaryotic transcription systems. A distinguishing feature of transcription initiation by Pol II, with respect to Pol I and Pol III, is that the latter two machineries can open promoter DNA spontaneously^36–40^, whereas the Pol II machinery generally requires the help of an ATP-dependent translocase subunit in TFIIH and its accessory factor TFIIE^22,41^. However, we show here that on snRNA promoters, mammalian Pol II, together with the factors that form the core PIC, can open DNA spontaneously without the help of TFIIE and TFIIH. Such spontaneous DNA opening has also been observed for yeast Pol II at a subset of promoters^31^ and also in the related archaeal transcription system^42^. Whereas spontaneous DNA opening occurs in the upstream-to-downstream direction, TFIIH-assisted DNA opening occurs in the downstream-to-upstream direction^22,41^. Our work thus provides evidence that, depending on the promoter, Pol II can use both types of DNA opening mechanisms, and suggests that TFIIH-assisted DNA opening originated later in the evolution of cellular DNA-dependent RNA polymerase machineries.

Several open questions remain to be addressed for a better understanding of snRNA gene transcription. In particular, SNAPc has been identified to be regulated by its direct interaction with activators that localize ~200 bp upstream of the PSE at the so-called distal sequence element (DSE)^7^. The intervening genomic region between PSE and DSE may be decorated by a nucleosome^8^. In the future, our work may be expanded to studying how DSE binding activators interact with the SNAPc-containing Pol II PIC described here, and how a nucleosome may enable or modulate this interaction. Additionally, our work also serves as stepping stone towards addressing the function of SNAPc in U6 snRNA transcription by Pol III. Such work should provide insights into how SNAPc can interact with both, the Pol II and the Pol III initiation machineries, providing further insights into the evolution of eukaryotic transcription systems.

## ONLINE METHODS

### Cloning and protein expression

cDNA constructs of SNAPc-FL containing SNAPC4 with an N-terminal StrepTwin-tag and a C-terminal His-tag, SNAPC1, SNAPC2, SNAPC3 and SNAPC5 were subcloned into the pLIB vector. The genes were assembled into a pBIG2ab vector using the biGBac system^43^. The cloned construct was transformed into DH10 EMBacY cells to generate bacmids. Next, the purified bacmid was mixed with CelfectinTM II reagent (Thermo Fisher Scientific) and transfected into 2 ml (density: 0.5 million cells/ml) of adherent Sf9 cells in a 6 well plate. After incubating the plate at 27 °C for 72 h, the resulting supernatant (P1 virus) was collected. To amplify the viral stock, 2 ml of P1 virus was added to 25 ml of Sf9 cells (0.5 million cells/ml) and incubated at 27 °C with shaking at 130 rpm. The supernatant (P2 virus) was collected after 4-5 days of infection when the cell viability dropped to <85% and stored at 4 °C. Large scale protein expression was carried out using 3 × 400 ml of High5 cells (0.5 million cells/ml) by adding 2 ml of P2 virus in each flask and incubated at 27 °C for 4 days at 130 rpm. Cells were then harvested by centrifugation at 250 × g for 10 mins at 4 °C, and pellets were stored at −80 ° C. SNAPc-core (SNAPC4 1-516 and lack of the SNAPC2 subunit) was expressed as previously described^18^.

### Protein purification

The insect cells pellet of SNAPc-FL were resuspended in buffer A containing 50 mM HEPES pH 7.8, 750 mM NaCl, 10% glycerol, 15 mM imidazole, 10 mM β-mercaptoethanol, 2 mM MgCl_2_, 1 mM phenylmethylsulfonyl fluoride (PMSF), 1 μg/mL Aprotinin, 1 μg/mL Pepstatin, and 1 μg/mL Leupeptin, supplemented with four EDTA-free protease inhibitor tablets (Pierce), a scoop of DNAse I, and 10 μl benzonase. Lysis was performed using a dounce homogeniser followed by sonication and the lysate was clarified by centrifugation at 48,000 × *g* at 4 °C for 40 mins. The supernatant was filtered using a 0.45-μm filter, and applied onto a HisTrap HP 5 ml column (GE Healthcare), pre-equilibrated with buffer A. The column was washed with 10 CV of buffer A1 (50 mM HEPES pH 7.8, 500 mM NaCl, 10% glycerol, 50 mM imidazole, 10 mM β-mercaptoethanol, 0.5 mM PMSF and 10 mM O-Phospho-L-serine), and then with 5 CV of buffer A2 (50 mM HEPES pH 7.8, 1250 mM NaCl, 10% glycerol, 50 mM imidazole, 10 mM β-mercaptoethanol, and 0.5 mM PMSF). The column was again washed with 5 CV buffer A1, and the bound protein complex was eluted in buffer B (50 mM HEPES pH 7.8, 500 mM NaCl, 10% glycerol, 300 mM imidazole, 10 mM β-mercaptoethanol, and 0.5 mM PMSF). Next, the sample was diluted to 250 mM NaCl with buffer Heparin A (50 mM HEPES pH 7.8, 10% glycerol, 1 mM TCEP, and 0.1 mM PMSF). The sample was centrifuged at 13,000 rpm for 15 mins at 4 °C and loaded onto a HiTrap Heparin HP 5 ml column (GE healthcare), pre-equilibrated with 12.5% of buffer Heparin B (50 mM HEPES pH 8, 2 M NaCl, 10% glycerol, 1 mM TCEP, and 0.1 mM PMSF). After washing with 5 CV of 12.5% buffer Heparin B, elution was performed through a linear gradient from 15% to 60% over 10 CV. The eluted fractions were analysed by SDS-PAGE, and fractions containing the SNAPc-FL complex were pooled, and concentrated using a 100 kDa molecular weight cut-off (MWCO) VivaSpin concentrator (Sartorius). The sample was centrifuged at 13,000 rpm for 15 mins at 4 °C and applied onto a Superose 6 PG XK 16/70 column (GE Healthcare), pre-equilibrated with 50 mM HEPES pH 7.8, 250 mM KCl, 10% glycerol, and 1 mM TCEP. Peak fractions were pooled, concentrated, flash-frozen and stored at −80 °C.

SNAPc-core has been purified as previously described^18^ with some modifications. Briefly, after cell lysis and centrifugation, the supernatant was subjected to nickel column purification (GE Healthcare) and eluted with 300 mM imidazole. The elution was then further purified with an heparin column and eluted with a gradient from 250mM to 1.25M NaCl. The fractions of interest were pooled, concentrated and subjected to size exclusion chromatography with a S200 16/600 equilibrated with 100 mM NaCl, 50 mM HEPES pH 7.9, 10% glycerol and 1mM TCEP. S. *scrofa* Pol II and human initiation factors TBP, TFIIA, TFIIB, TFIIE, TFIIF and TFIIH were purified as previously described^22^.

### Electrophoretic Mobility Shift Assays

EMSA was performed using a 76 bp fragment of U1 promoter DNA (template:5’-GAA ACG TTG TGC CTC TGC CCC GAC ACA GCC TCA TAC GCC TCA CTC TTT ACA CAC ACG GTC ACT TG CCC CGC GCA CT-3’ and its complementary strand) and a 75 bp fragment of U5 promoter DNA (template:5’-ACC AGT TAC TTC TGT AAC TCA ATT TTC GGG TAA CTG CAA TTC CTA GTA CAC TGA TGG TGT CTA CTA ATC CC AAG G-3’ and its complementary strand; Integrated DNA Technologies). 20 pM of SNAPc FL or core were incubated with 5 pM of annealed oligonucleotides in presence or absence of 25 pM of TFIIB and TBP in 20 μL of incubation buffer (250 mM NaCl, 50 mM HEPES pH 7.9, 20% glycerol, 1 mM TCEP) at room temperature for 15 min. The complexes were resolved on 5% polyacrylamide (37.5:1 acrylamide/bisacrylamide, 10% glycerol, Tris Borate EDTA 1x) gels in 0.5X Tris Borate EDTA running buffer at 40 mA. After staining with Ethidium bromide, the gels were scanned with a Typhoon FLA9500 (GE Healthcare).

### Promoter-dependent *in vitro* transcription assay

*In vitro* transcription assays were performed as described previously^22,41^ with minor alterations. The DNA scaffold (dsDNA) was prepared as reported using a pUC119 vector into which a 92 nucleotide fragment of the native U1 snRNA promoter^20^ had been inserted. The scaffold (non-template: 5’-GGG CGT GAC CGT GTG TGT AAA GAG TGA GGC GTA TGA GGC TGT GTC GGG GCA GAG GCA CAA CGT TTC GCC CGA AGA TCT CAT ACT TAC CTG GCA GGG CTA AGC TTG GCG TAA TCA TGG TCA TAG CTG TTT CCT GTG TGA AAT TGT TAT CCG CTC ACA ATT CCG CCC-3’, template: 5’-GGG CGG AAT TGT GAG CGG ATA ACA ATT TCA CAC AGG AAA CAG CTA TGA CCA TGA TTA CGC CAA GCT TAG CCC TGC CAG GTA AGT ATG AGA TCT TCG GGC GAA ACG TTG TGC CTC TGC CCC GAC ACA GCC TCA TAC GCC TCA CTC TTT ACA CAC ACG GTC ACG CCC-3’) was stored in low salt buffer (60 mM KCl, 10 mM K-HEPES pH 7.5, 8 mM MgCl_2_, 3% (v/v) glycerol).

Initiation complexes for *in vitro* transcription were reconstituted on scaffold DNA essentially as described^22,41^. All incubation steps were performed at 25 °C unless indicated otherwise. Per sample, 1.6 pmol scaffold, 1.8 pmol Pol II, TFIIE and TFIIH, 5 pmol TBP and TFIIB, 9 pmol TFIIF and TFIIA and 5 pmol SNAPc-FL or SNAPc-core were used. SNAPc was mixed and added to the sample simultaneously with TFIIB. Reactions were prepared in a sample volume of 23.8 μl with final assay conditions of 60 mM KCl, 3 mM K-HEPES pH 7.9, 20 mM Tris-HCl pH 7.9, 8 mM MgCl_2_, 2% (w/v) PVA, 3% (v/v) glycerol, 0.5 mM 1,4- dithiothreitol, 0.5 mg ml^−1^ BSA and 20 units RNase inhibitor. To achieve complete PIC formation, samples were incubated for 45 min at 30 °C. Transcription was started by adding 1.2 μl of 10 mM NTP solution and permitted to proceed for 60 min at 30 °C. Reactions were quenched with 100 μl Stop buffer (300 mM NaCl, 10 mM Tris-HCl pH 7.5, 0.5 mM EDTA) and 14 μl 10% SDS, followed by treatment with 4 μg proteinase K (New England Biolabs) for 30 min at 37 °C. RNA products were isolated from the samples as described^41^, applied to urea gels (7 M urea, 1x TBE, 6% acrylamide:bis-acrylamide 19:1) and separated by denaturing gel electrophoresis (urea-PAGE) in 1x TBE buffer for 45 minutes at 180 volts. Gels were stained for 30 min with SYBR™ Gold (Thermo Fisher Scientific) and RNA was visualized with a Typhoon 9500 FLA imager (GE Healthcare Life Sciences).

### Preparation of the SNAPc-containing Pol II PIC

We performed the assembly of SNAPc containing Pol II PIC on snRNA promoters at 25°C essentially as described previously. We used a 96bp fragment of both the native U1 promoter DNA(template:5’-ATC ATG GTA TCT CCC CTG CCA GGT AAG TAT GAA ACG TTG TGC CTC TGC CCC GAC ACA GCC TCA TAC GCC TCA CTC TTT ACA CAC ACGGTC ACT TGC-3’;non-template: 5’-GCA AGT GAC CGT GTG TGT AAA GAG TGA GGC GTA TGA GGC TGT GTC GGG GCA GAG GCA CAA CGT TTC ATA CTT ACC TGG CAG GGG AGA TAC CAT GAT-3’) and an engineered U5 promoter with 10bp deleted from the downstream edge of the PSE sequence (template: 5’-CCC TGC CAG GTT TTA TGC GAT CTG AAG AGA AAC CAG AGT ATA CCA GTT ACT TCT GTA ACT CAA TTT TCG GGT CCTAGT ACA CTG ATG GTG TCT ACT-3’; non-template: 5’-AGT AGA CAC CAT CAG TGT ACT AGG ACC CGA AAA TTG AGT TAC AGA AGT AAC TGG TAT ACT CTG GTT TCT CTT CAG ATC GCA TAA AAC CTG GCA GGG-3’). In summary, SNAPc (FL or Core) was pre-incubated for 5 min with the snRNA promoter (U1 or U5) scaffold. It was then mixed with TFIIA-TFIIB and TBP followed by the pre-formed Pol II-TFIIF complex. TFIIE was then added to this mixture and the assembly was incubated at 25°C for 60 min at 300 rpm. This reconstituted SNAPc containing Pol II PIC was subjected to 10-30% sucrose-gradient ultra-centrifugation with simultaneous cross-linking using GraFix (Kastner et al., 2008) at 175,000*g* for 16h at 4°C. The assay was then fractionated as 200μl aliquots where the crosslinking reaction was quenched using a cocktail of 10mM aspartate and 30mM lysine for 10mins. Fractions with SNAPc containing Pol II PIC were dialysed against the cryo-EM sample buffer (25 mM HEPES pH 7.6, 100 mM KCl, 5 mM MgCl_2_, 1% glycerol and 3 mM TCEP).

### Cryo-EM data collection and processing

Samples for cryo-EM were prepared using Quantifoil R3.5/1 holey carbon grids pre-coated with a homemade 3 nm continuous carbon. Four microlitres of SNAPc containing Pol II PIC sample bound to snRNA promoter (U1/U5) was added to the carbon side and incubated for 2.5 min. The grids were blotted for 2.5 s and vitrified by plunging into liquid ethane with a Vitrobot Mark IV (FEI Company) set at 4 °C and 100% humidity. Cryo-EM data were collected on a 300-kV FEI Titan Krios with a K3 summit direct detector (Gatan) and a GIF quantum energy filter (Gatan) operated with a slit width of 20 eV. Automated data collection was performed with SerialEM at a nominal magnification of 81,000x, corresponding to a pixel size of 1.05 Å/pixel^44^. For the sample containing U1 promoter, a total of 16,854 image stacks, with each stack containing 50 frames, were collected at a defocus range of −0.5 to −3.0 μm. All movie frames were contrast transfer function (CTF)-estimated, motion-corrected and dose-weighted using Warp^45^. Particles were picked by Warp using a trained neural network, resulting in 5,181,947 particles as a starting set. Subsequent steps of image processing were performed with cryoSPARC^46^ and RELION v.3.1.0^47^.

Particles were extracted with a binning factor of 2 and a box size of 200 pixels (a pixel size of 2.1 Å/pixel) to perform initial clean-up and sorting. The processing scheme was centered around identifying the best SNAPc-containing particle sets. Iterative rounds of 2D-classification followed by heterogenous and homogenous refinements in cryoSPARC, led to two sets of particles corresponding to CC (set-1: 252,067 particles) and OC (set-2: 240,243 particles) promoter states respectively. Each set was re-extracted without binning and processed using RELION v.3.1.0, as follows. For set-1, the particles were further sorted by focused 3D classification with a large spherical mask (Mask-1) encompassing the upstream region of PIC containing SNAPc, TBP, TFIIA and TFIIB. This resulted in identifying the best 47,293 SNAPc-containing particles. These particles were again subjected to 3D refinement using Mask-1, giving rise to a reconstruction of SNAPc containing Pol II PIC bound to U1 promoter in CC state at 3.4 Å resolution (map-1). In parallel, focused 3D classification of set-2 with a spherical mask (Mask-2) around the upstream region helped to identify the best 137,246 SNAPc containing particles. These particles were then subjected to 3D refinement followed by CTF refinement and Bayesian polishing. Following this, the particles were subject to refinement with and without mask-1 to obtain of SNAPc containing Pol II PIC bound to U1 promoter in OC state at 3.0 Å (map-2) and a local map spanning the SNAPc containing upstream region at 3.7 Å resolution(map-3).

For the sample containing U5 promoter dataset, 4,842image stacks, with each stack containing 60 frames, were collected at a defocus range of −0.3 to −2.5 μm. All movie frames were contrast transfer function (CTF)-estimated, motion-corrected and dose-weighted using Warp^45^. Particles were picked by Warp using a trained neural network, resulting in 1,299,523 particles. Subsequent steps of image processing were performed with cryoSPARC^46^ and RELION v.3.1.0^47^. Particles were extracted with a binning factor of 4 and a box size of 100 pixels (a pixel size of 4.2 Å/pixel) to perform initial clean-up and sorting. After sorting in cryoSPARC using 2D-classification followed by heterogenous and homogenous refinements, a particle set (set-3: 443,960 particles) in CC promoter state was re-extracted with 2x binning (a pixel size of 2.1 Å/pixel) and processed using RELION v.3.1.0, as follows. For set-3, the particles were further sorted by 3D classification followed by focused 3D classification using Mask-1. The resulting 159,144 particles were re-extracted without a binning factor and were subjected to CTF refinement and Bayesian polishing. These particles were then subjected to another round of masked classification yielding 85,787 SNAPc-containing particles. These particles were then 3D refined without and with Mask-1, giving rise to a reconstruction of SNAPc containing Pol II PIC bound to U5 promoter in CC state at 3.0 Å resolution (map-4) and a local map of the SNAPc containing upstream complex extending to 3.2 Å(map-5).

The reported resolutions were calculated on the basis of the gold standard Fourier shell correlation (FSC) 0.143 criterion. After processing of the final reconstructions, B-factor sharpening was performed for all final maps on the basis of automatic B-factor determination in RELION (−5 Å^2^ for map-1: SNAPc-PIC bound to U1 promoter in CC state, −10 Å^2^ for map-2: SNAPc-PIC bound to U1 promoter in OC state and −10 Å^2^ for map-3: local map of SNAPc containing upstream complex, −10 Å^2^ for map-4: SNAPc-PIC bound to U5 promoter in CC state and −10 Å^2^ for map-5: local map of SNAPc containing upstream complex). Estimates of local resolution were calculated using the in-built local-resolution tool of RELION and the estimated B-factors. To assist model building, a local-resolution-filtered map (but unsharpened) of map-5 was sharpened locally using PHENIX.auto_sharpen^48^.

### Model building and refinement

The PIC was modelled using the core PIC part of the previously published high resolution structures in closed and open promoter states^22^. For SNAPc, the subunits SNAPC1 and SNAPC4 were built using partial homology models generated using TrRosetta^49^. The partial models were rigid body fitted into the density using UCSF Chimera^50^ and were manually extended and corrected using Coot^51^ to fit the density. The subunit SNAPC3 was modelled entirely de novo using the experimental density in Coot. Ambiguous density corresponding to linker regions were not modelled. The model corresponding to the wing-2 region constituting parts of SNAPC1, SNAPC3 and SNAPC4 was modelled using AlphaFold^27^. The model for promoter DNA in CC and OC states was obtained using the high-resolution structures of human PIC as template where in the sequence register was mutated to fit the U1 and U5 respectively. The models were then subjected to iterative rounds of PHENIX real-space refinement followed by manual adjustment in coot to achieve final models with good stereochemistry as assessed by MolProbity^52^. Figures representing the 3D structures and maps were prepared using PyMOL, UCSF Chimera and UCSF ChimeraX.

### Crosslinking mass-spectrometry

To prepare a sample for performing crosslinking mass-spectrometry, a stable complex of SNAPc-containing Pol-II PIC bound to U5 promoter was isolated. An assay containing Pol II, TBP, TFIIA, TFIIB, TFIIF and SNAPc-FL was incubated in ratios explained above and was subjected to size-exclusion chromatography using Superose 6 increase 3.2/300 GL column (GE Healthcare) pre-equilibrated with buffer-x (25mM Hepes pH 7.5, 100mM NaCl, 5mM MgCl2, 5% glycerol and 2mM TCEP). The peak fractions were then pooled and incubated with 1mM of Bissulfosuccinimidyl suberate (BS3) for 45 min at 4° C. The crosslinking reaction was quenched using a cocktail of 10 mM aspartate and 30 mM lysine.

Crosslinked proteins were resuspended in 4 M urea/ 50 mM ammonium bicarbonate for 10 min at 25°C and reduced for 30 min at RT with 10 mM dithiotreitol (DTT). Proteins were alkylated for 30 min at RT in the dark by adding iodacetamide (IAA) to a final concentration of 55 mM. Sample was diluted to 1M Urea and digested for 30 min at 37 °C with 4 μl Pierce Universal Nuclease (250 U/μl) in the presence of 2 mM MgCl2. Trypsin (Promega) digest was performed o/n at 37 °C in a 1:50 enzyme/protein ratio, the reaction was terminated with 0.2 % (v/v) FA. Tryptic peptides were desalted on MicroSpin Columns (Harvard Apparatus) following manufacturer’s instruction and vacuum-dried. Cross-linked peptides were resuspended in 50 μl 30 % acetonitrile/0.1 % TFA and enriched by peptide size exclusion chromatography/pSEC (Superdex Peptide PC3.2/300 column, GE Healthcare, flow rate 50 μl/min).

Crosslinked peptides derived from pSEC were subjected to liquid chromatography mass spectrometry (LC-MS) on a Thermo Obitrap Exploris mass spectrometer. Peptides were loaded in duplicates onto a Dionex Ultimate 3000 RSLCnano equipped with a custom column (ReproSil-Pur 120 C18-AQ, 1.9 μm pore size, 75 μm inner diameter, 30 cm length, Dr. Maisch GmbH). Peptides were separated applying the following gradient: mobile phase A consisted of 0.1 % formic acid (FA, v/v), mobile phase B of 80 % ACN/0.08 % FA (v/v). The gradient started at 5 % B, increasing to 10, 15 or 20 % B within 3 min, followed by a continuous increase to 48 % B within 45 min, then keeping B constant at 90 % for 8 min. After each gradient, the column was again equilibrated to 5 % B for 2 min. The flow rate was set to 300 nL/min.

MS1 spectra were acquired with a resolution of 120,000 in the orbitrap (OT) covering a mass range of 380–1600 m/z. Dynamic exclusion was set to 30 s. Only precursors with a charge state of 3-8 were included. MS2 spectra were recorded with a resolution of 30,000 in OT and the isolation window to 1.6 m/z. Fragmentation was enforced by higher-energy collisional dissociation (HCD) at 30 %. Raw files were searched against a database containing the sequences of the proteins of the complex and analyzed via pLink 2.3.9 at a false discovery rate (FDR) of 1%^53^. Carbamidomethylation of cysteines was set as fixed modification, oxidation of methionines as variable modification. The database contained all proteins within the complex. For further analysis only interaction sites with 3 cross-linked peptide spectrum matches were taken into account. Cross-links were displayed with xiNET and XlinkAnalyzer in UCSF Chimera.^50,54,55^

### TSS precision analyses in cells

We utilized published 5’cap-seq data^35^ (GEO: GSE159633) for analyses of TSS precision in cells. The raw data were processed as described previously^35^ to obtain the 5’-ends of reads and generate normalized coverage. In brief, we first removed the unique molecular identifier (UMI) from 5cap-seq reads with UMI-tools^56^ and then trimmed adapter sequences with Cutadapt ^57^ and mapped to the human genome (GRCh38) merged with the D. melanogaster genome (Dm6) with the STAR mapper^58^. We next deduplicated the mapped data with UMI-tools to remove any PCR duplicates and then determined the first transcribed base and used this position in downstream analyses. Normalization factors were obtained from the spike-in reads (processed as above) that mapped to the spike-in genome and used to normalize the human genome coverage profiles. The replicates were combined by summing the normalized coverage per nt. Thus, obtaining genome-wide capped 5’-end signal (5’cap-seq signal) at single-base resolution. We subset the NCBI reference genome annotation^59^ (GRCh38.p7) to only contain genes annotated to the primary assembly and included only genes with known transcripts (prefix: “NR” or “NM”) and also excluded overlapping genes. To exclude genes with alternative start sites from downstream analyses we included only genes that have a constitutive first or a single exon in our downstream analyses.

To determine the main TSS we determined the position with the highest 5’cap-seq signal within constitutive first exons of the reference annotation. To accommodate for reference annotation imprecision, we also included 10 bp upstream of the annotated TSS and set the downstream cutoff to 500 bp downstream of the annotated TSS. We thus obtained the main TSS for each constitutive TSS. We next quantified the 5’cap-seq signal of the main TSS (±2 bp) and the TSS region (main TSS ±50 bp). We excluded genes with less than 10 counts in the TSS region and genes with biotypes that are not either protein-coding or snRNA. From the remaining annotated snRNA subset we also removed know Pol III-transcripts: RN7SK, RNU6ATAC, SNAR-G2, RNU6-2, SNAR-C4, SNAR-G1, SNAR-C3 and identified with protein-coding gene promoters contain a TATA-box motif (JASPAR database, 2020 release: https://jaspar2020.genereg.net/matrix/POL012.1/) within 50 bp upstream of the annotated TSS. Finally, we determined the TSS precision score by dividing the TSS peak counts by the TSS region counts. The maximum TSS precision score is 1, which means that all 5’cap-seq signal is within the TSS peak. The preprocessed 5’cap-seq data was analyzed in RStudio^60^ utilizing R version 3.6.1^61^ and packages from the Bioconductor repository ^62,63^ and Tidyverse^64^. Plots were generated with ggplot2 and ggbio ^65^.

## Acknowledgements

We thank present and past members of the Cramer laboratory for help and discussions. We thank Frauke Grabbe for purifying TBP, TFIIA, TFIIB, TFIIE, TFIIF and TFIIH. We thank C. Dienemann and S. Schilbach for input towards designing the engineered U5 promoter DNA. We thank C. Dienemann and U. Steuerwald for maintenance of the electron microscopy facility. S.R. was supported by a postdoctoral fellowship from Peter und Traudl Engelhorn Foundation. A.V. was supported by the Cancer Research UK Programme Foundation (CR-UK C47547/A21536) and a Wellcome Trust Investigator Award (200818/Z/16/Z. P.C. was supported by the Deutsche Forschungsgemeinschaft (EXC 2067/1 39072994, SFB860) and the ERC Advanced Investigator Grant CHROMATRANS (grant agreement No. 882357).

## Author Contributions

S.R. carried out all experiments and data analysis, unless stated otherwise. S.S. performed the *in vitro* transcription assay and quantification. T.K. and J.G. cloned, expressed and purified the SNAPc variants and performed EMSA assays. K.Z. performed the reanalysis of 5’-capseq data, TSS precision plots and the web-logo plots. J.S. performed crosslinking mass-spectrometry and data analysis and was supervised by H.U. S.R. and C.D. collected the cryo-EM datasets. P.C. and A.V. designed and supervised research. S.R. and P.C. interpreted the data and wrote the manuscript, with input from all authors.

## Competing interests

The authors declare no competing interests.

**Supplementary Information** is available online at https://doi.org/XYZ.

## Correspondence

Correspondence and request of materials and resources should be addressed to P.C.

## Data availability

The cryo-EM density reconstructions were deposited to the EMDB under accession codes EMD-AAAA, -BBBB, -CCCC, -DDDD, -EEEE and atomic coordinates were deposited to the PDB under the accession codes PDB-AAAA, -BBBB, -CCCC, -DDDD, -EEEE. All data is available in the main text or the supplementary materials.

## EXTENDED DATA FIGURES

**Extended Data Figure 1.**
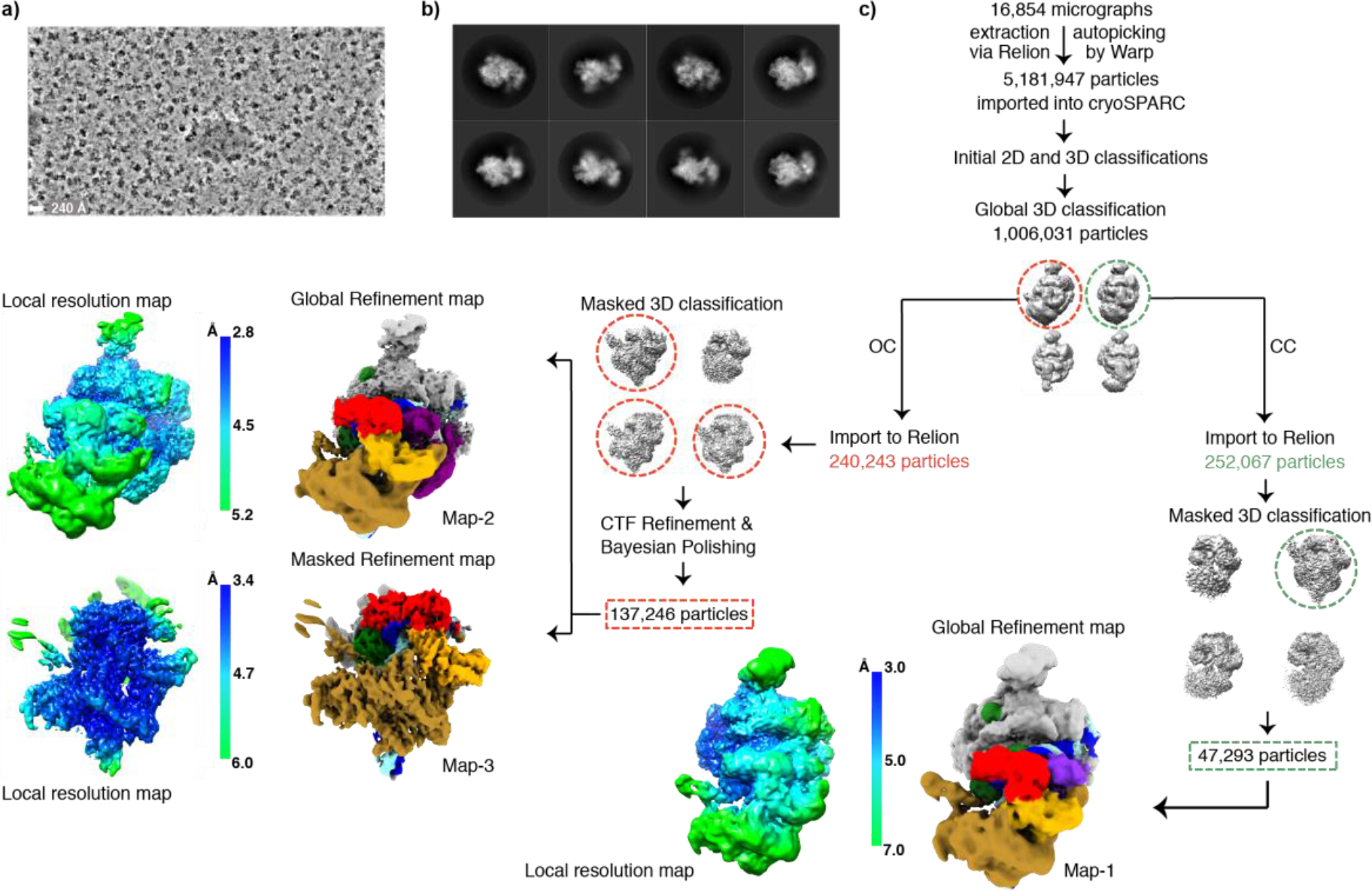
Processing of cryo-EM data for SNAPc-containing Pol II PIC bound to U1 promoter. Related to figure 2. a) Representative cryo-EM micrograph of the SNAPc-containing Pol II PIC bound to U1 promoter cryo-EM data collection. Scale bar – 240 Å b) Representative 2D class averages of initially sorted datasets after merging. Adjacent to a well-defined PIC, clear signal for SNAPc is detected. c) Complete processing scheme. After initial clean-up procedures, particles representing SNAPc containing PIC were recovered as two sets. These particle sets were processed separately with respect to the promoter DNA state (CC/OC) and SNAPc occupancy. Final maps are coloured using the subunit color code in Figure 1. The local resolution map indicate the resolution range of final maps (scale bar).

**Extended Data Figure 2.**
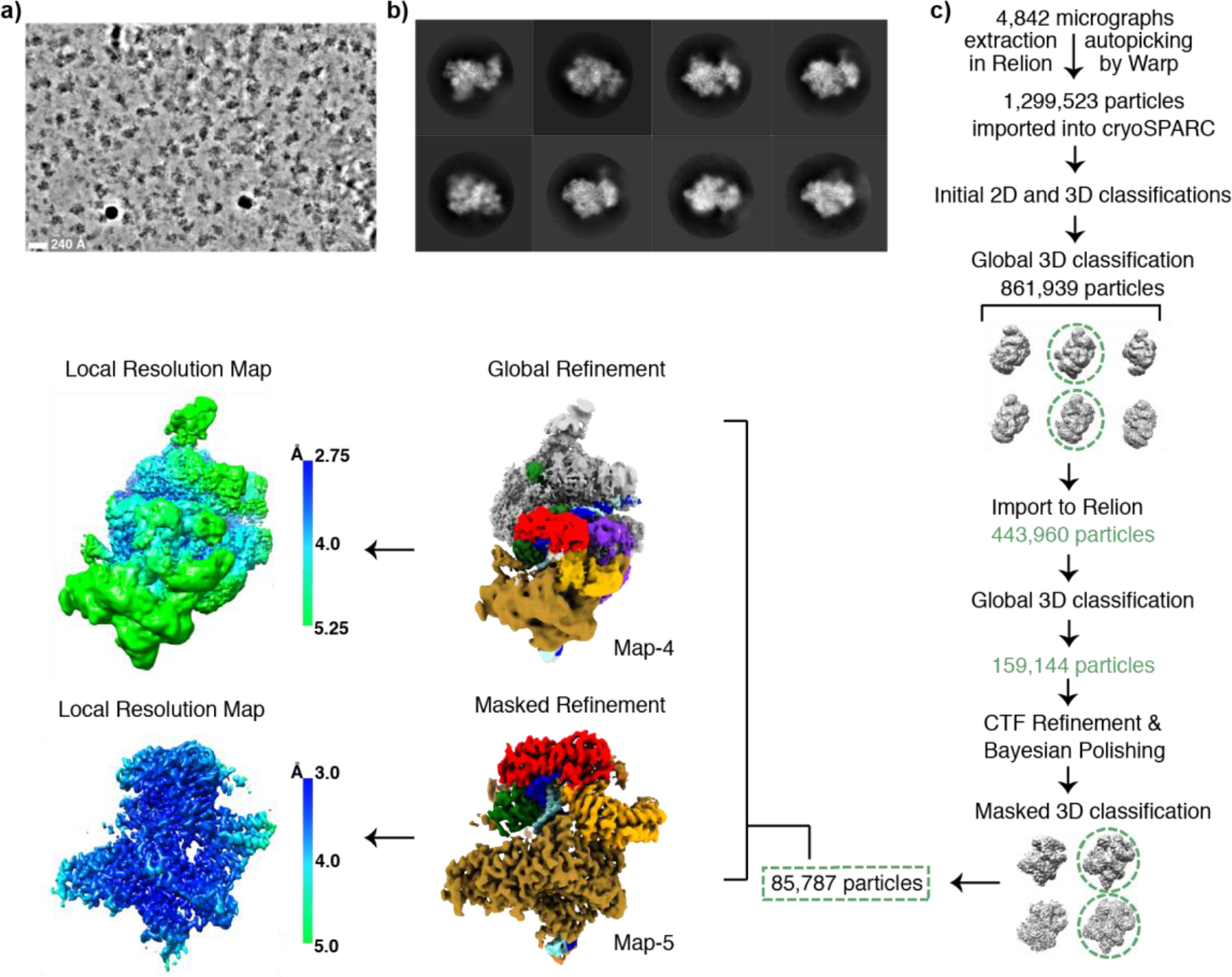
Processing of cryo-EM data for SNAPc-containing Pol II PIC bound to U5 promoter. Related to Figure 2. a) Representative cryo-EM micrograph of the SNAPc-containing Pol II PIC bound to U5 promoter cryo-EM data collection. Scale bar – 240 Å. b) Representative 2D class averages of initially sorted datasets after merging. As in the case of U1 promoter dataset, a clear signal for SNAPc is detected adjacent to a well-defined PIC. c) Complete processing scheme. The optimized strategy from U1 promoter bound SNAPc-PIC dataset was used to obtain high resolution maps of SNAPc-PIC bound to U5 promoter. Final maps are coloured using the subunit color code in Figure 1. The local resolution map of global and locally refined maps indicate the resolution range of final maps (scale bar).

**Extended Data Figure 3.**
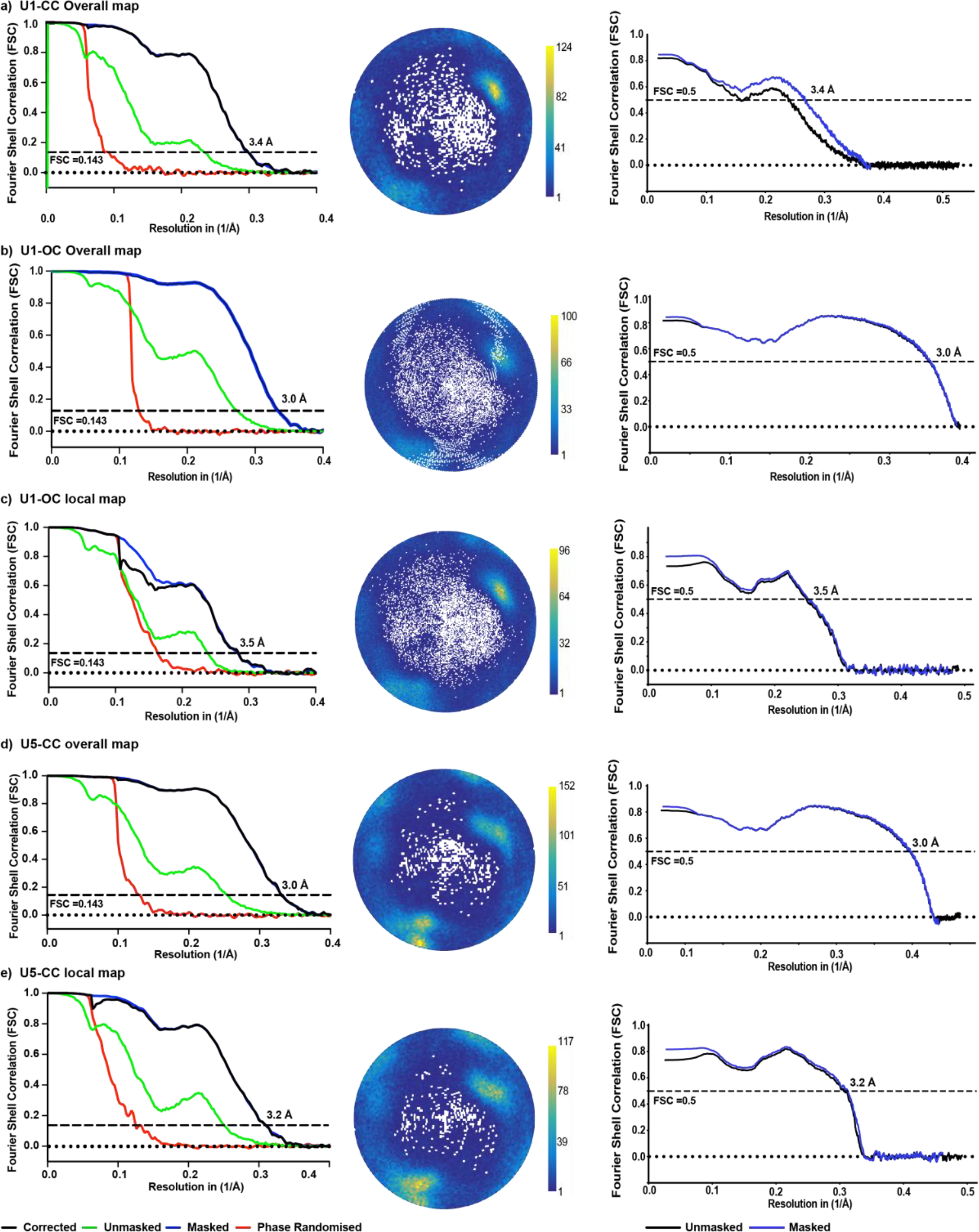
FSC and angular distribution plot of cryo-EM reconstructions. Related to Figure 2. a-e) On the left - FSC plot showing the overall resolution of the reconstructions determined by the gold standard FSC cut-off 0.143, indicated in the graph. In the middle – angular distribution plot of the respective reconstruction showing assignment of particles with respect to various angles. Colour bar indicates number of samples per angular bin (white areas indicate unpopulated angles). On the right - Model-to-map FSCs, showing the fit of modelled structures to their corresponding maps.

**Extended Data Figure 4.**
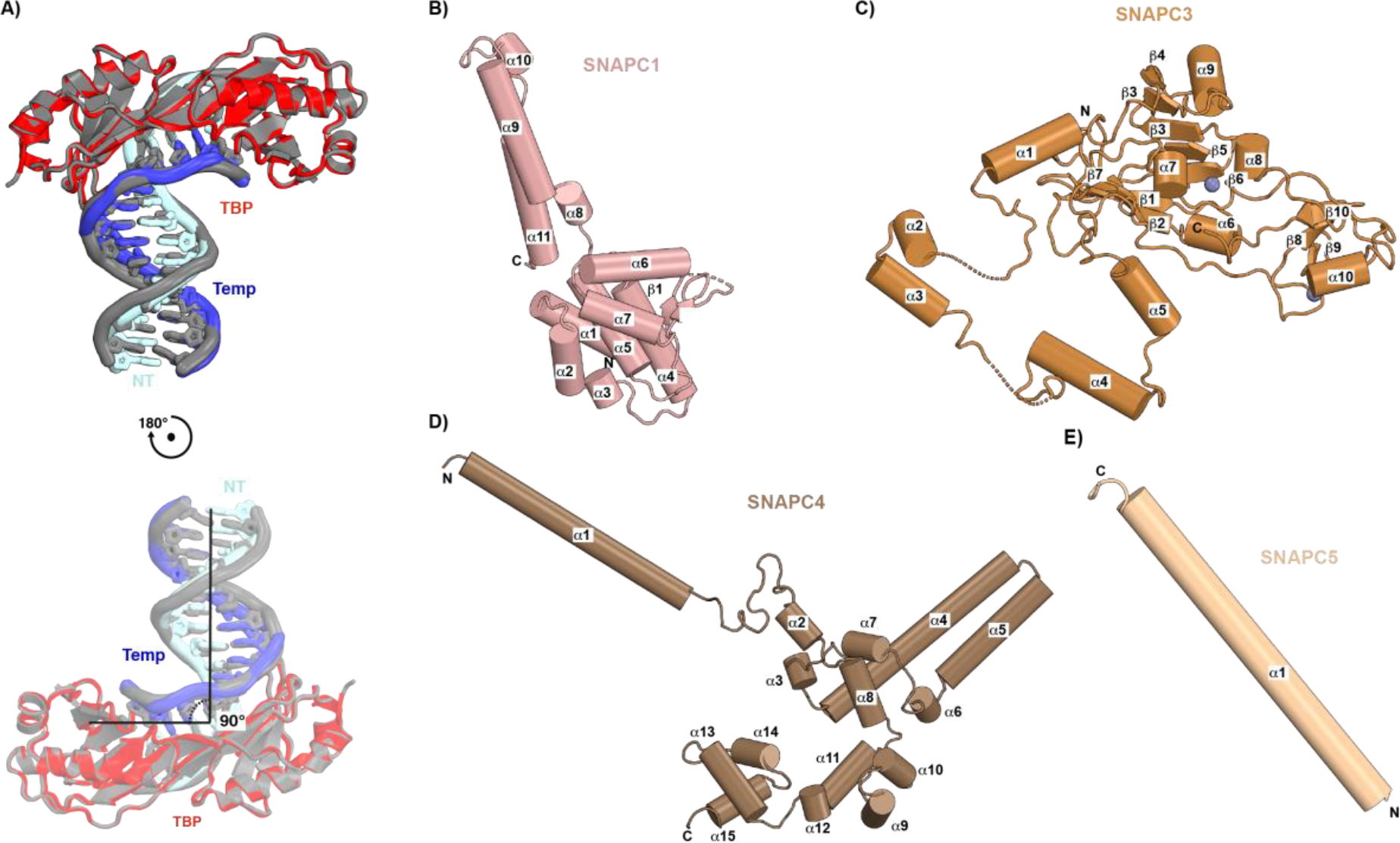
Structural comparison of TBP bound to TATA containing and TATA-less DNA template; Overall of Structure of individual SNAPc subunits. Related to Figures 2 and 3. a) Structural super-position of TBP(red) bound TATA-less U1 promoter (cyan/blue) on to TBP (grey) bound to TATA box sequence (PDB: 1YTF)(Tan et al., 1996). The comparison shows that TBP binds to the TATA-less sequence in a canonical fashion and bends the DNA by 90° b-e) Cartoon representation of the individual structures of SNAPc subunits SNAPC1, 3, 4 and 5 displaying its secondary structure elements as labelled. The N and C termini of all subunits are indicated.

**Extended Data Figure 5.**
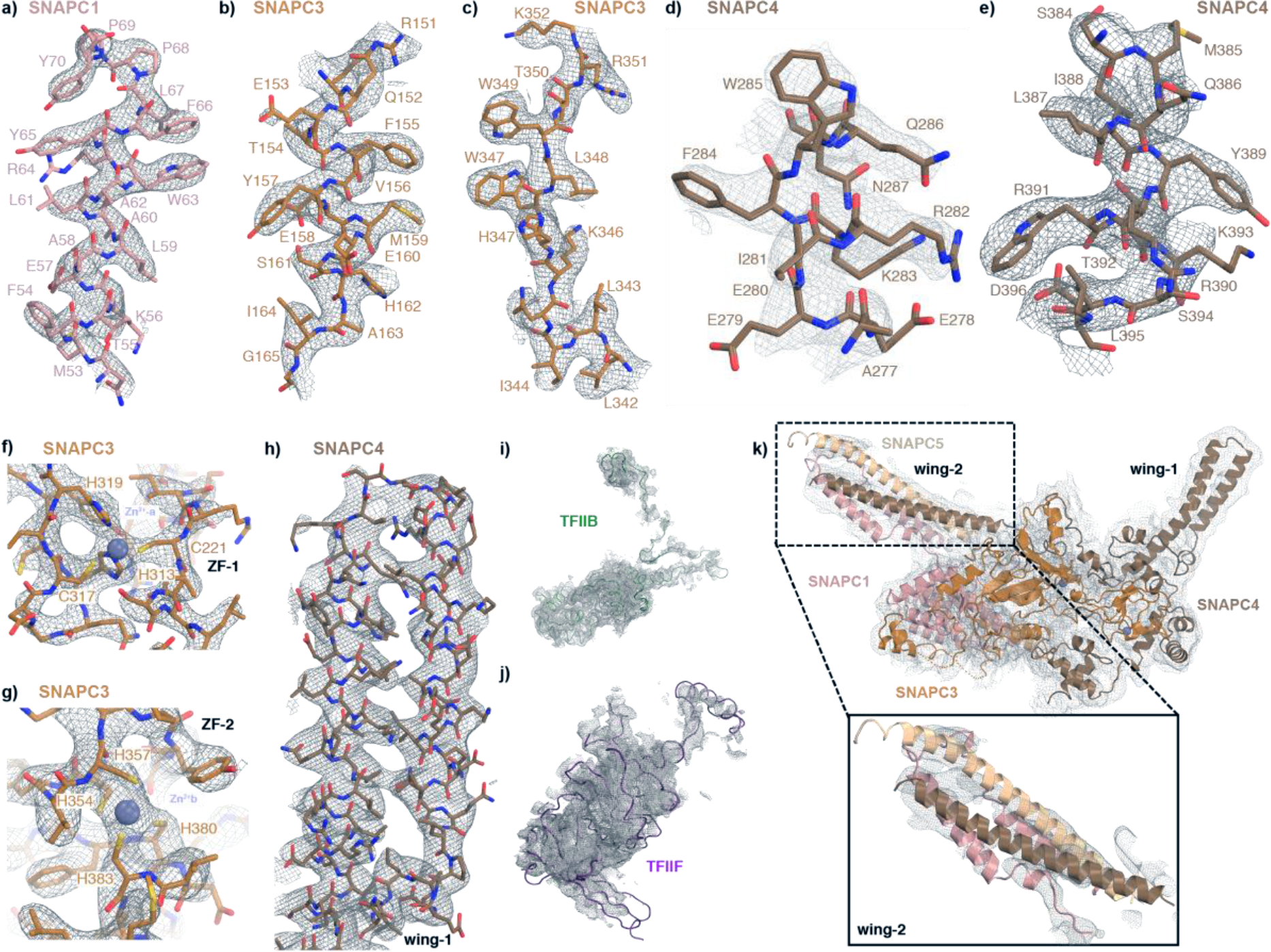
Map quality and map to model fit. Related to Figure 3. a-h) Sections of cryo-EM density of SNAPc subunits overlaid with their respective atomic models. Densities are shown as a grey mesh, and sticks are shown for the model as coloured in Figure 3. i) cryo-EM density of the TFIIB subunit overlaid to the atomic within the SNAPc containing Pol II PIC bound to U1 promoter in OC state. j) cryo-EM density of a region of TFIIF subunit overlaid to the atomic model within the SNAPc containing Pol II PIC bound to U1 promoter in OC state. d) Local map of SNAPc containing Pol II PIC bound to U1 promoter in OC state is low pass filtered to 5Å. The corresponding map is fitted with SNAPc subunits representing map to model fit, in particular the ‘wing-2’ region modelled using AlphaFold2^27^.

**Extended Data Figure 6:**
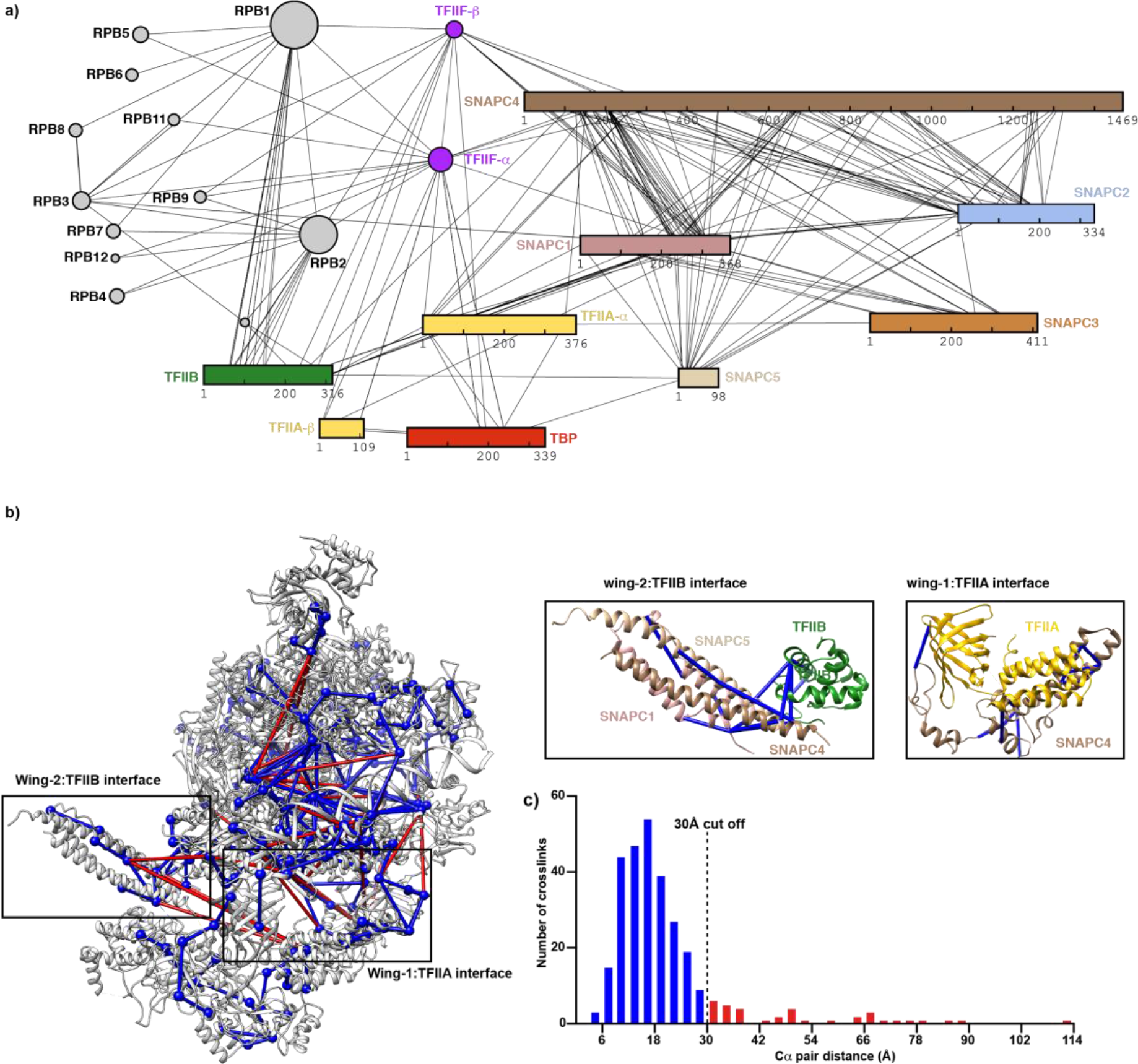
Crosslinking mass-spectrometric analysis of SNAPc containing Pol II PIC. Related to Figures 2 and 3. a) 2D representation of the overview of BS3 crosslinks. The crosslinks correspond to inter-protein mono-links that have at least three crosslinked peptide-spectrum matches (CSM). The subunit colours are consistent with Figure 2. b) Crosslinks as mapped to SNAPc containing Pol II PIC structure using Xlink analyzer^55^ plugin in UCSF chimera. The inset show the crosslinks observed between SNAPc subunits and the GTFs’ TFIIA and TFIIB respectively. c) Histogram representing the distribution of Cα pair distances of unique crosslinks mapped to the structure. Dotted line indicates the 30Å cut-off for BS3 crosslinked Cα pair. A total of 87.8% of the crosslinks were satisfied within this 30 Å cutoff.

**Extended Data Figure 7.**
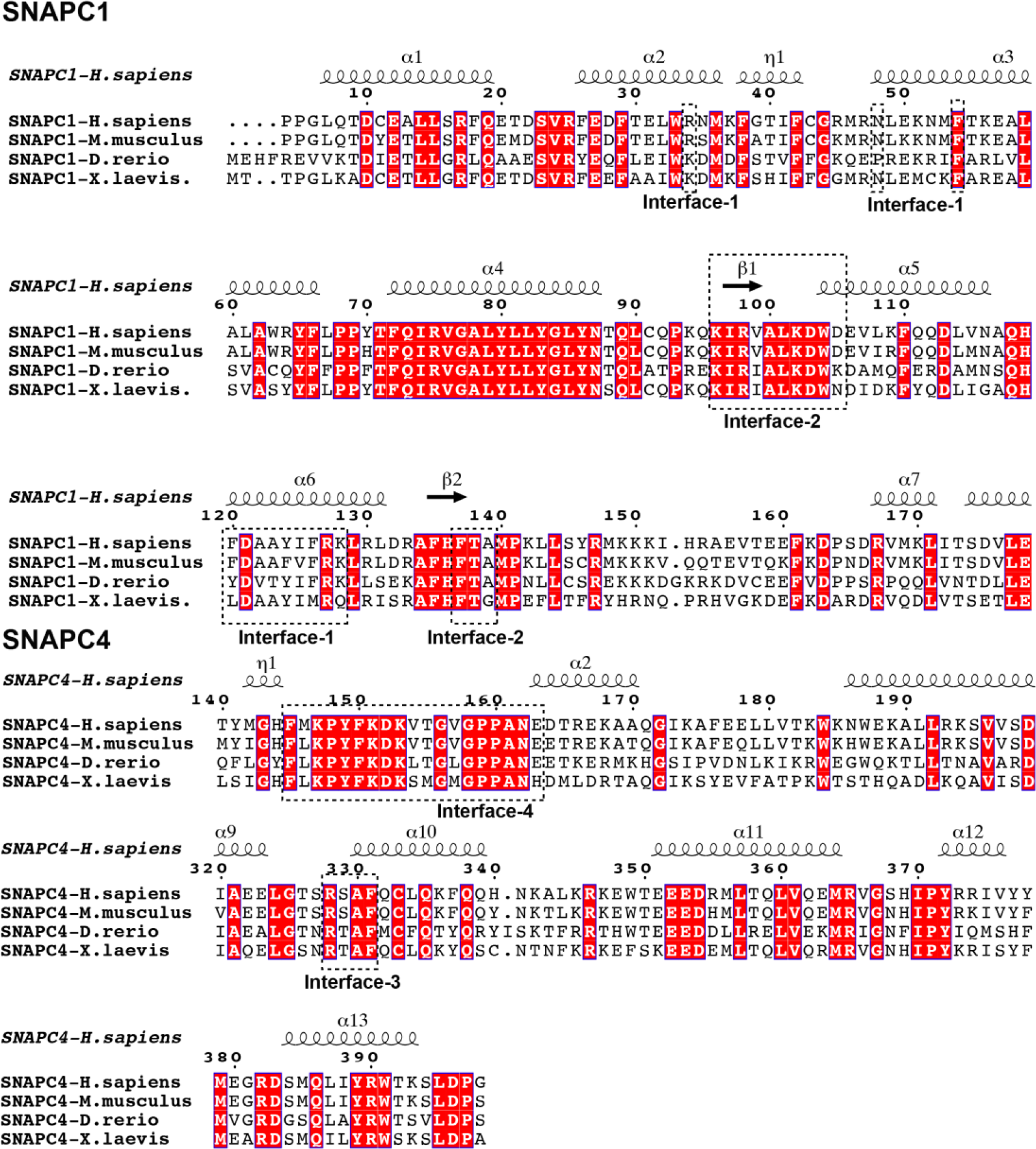
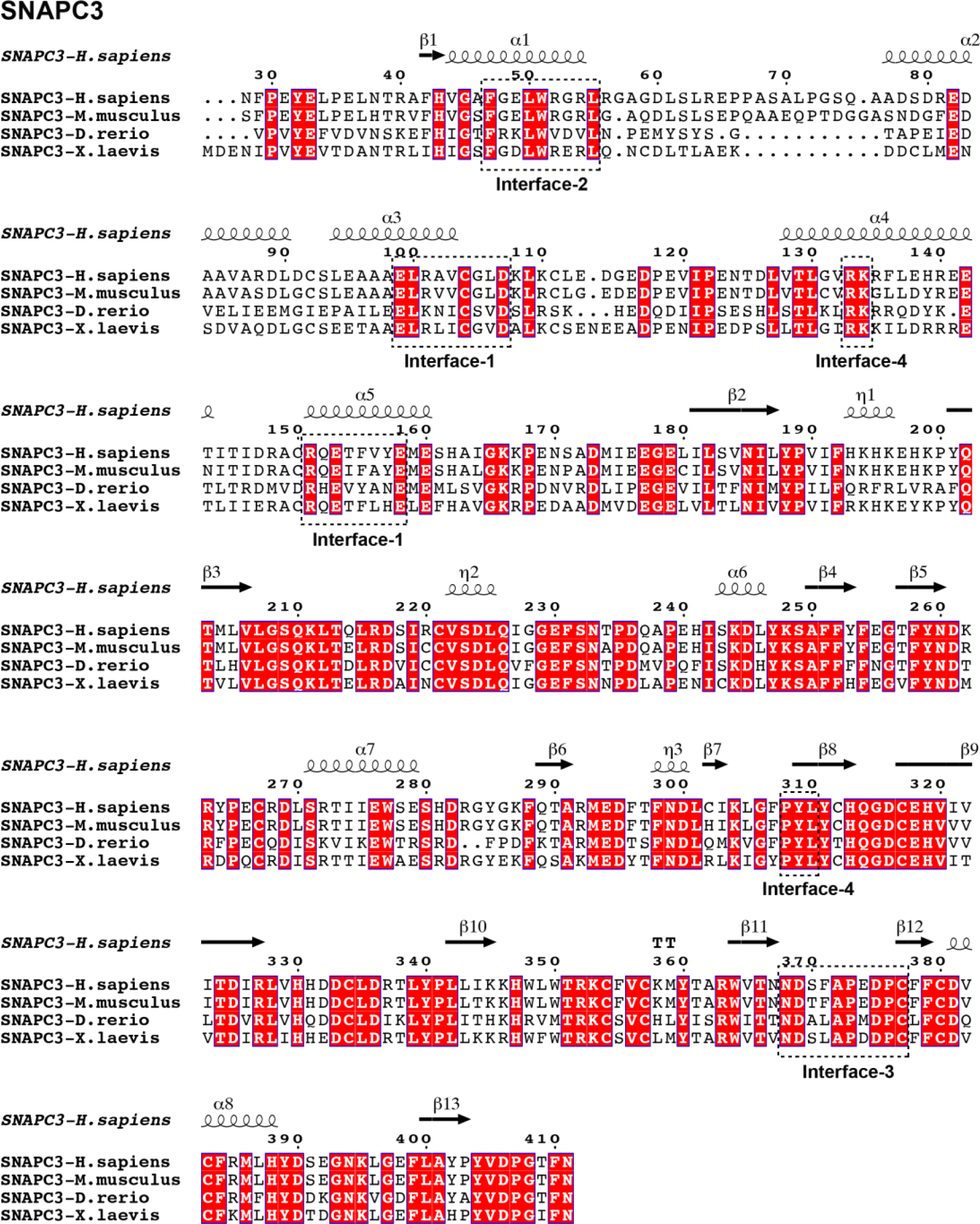
Structure based sequence alignment of SNAPc subunits involved in interactions. Related to Figures 3 and 4. Sequence alignments were performed with the regions of individual subunits for which the structure has been determined in this study. T-Coffee algorithm^66^ was adopted to obtain a structure based sequence alignment which was then visualized using ESPript^67^. Residues with identity above 80% are coloured red. Regions involved in interactions are indicated by dashed boxes and labels.

**Extended Data Figure 8:**
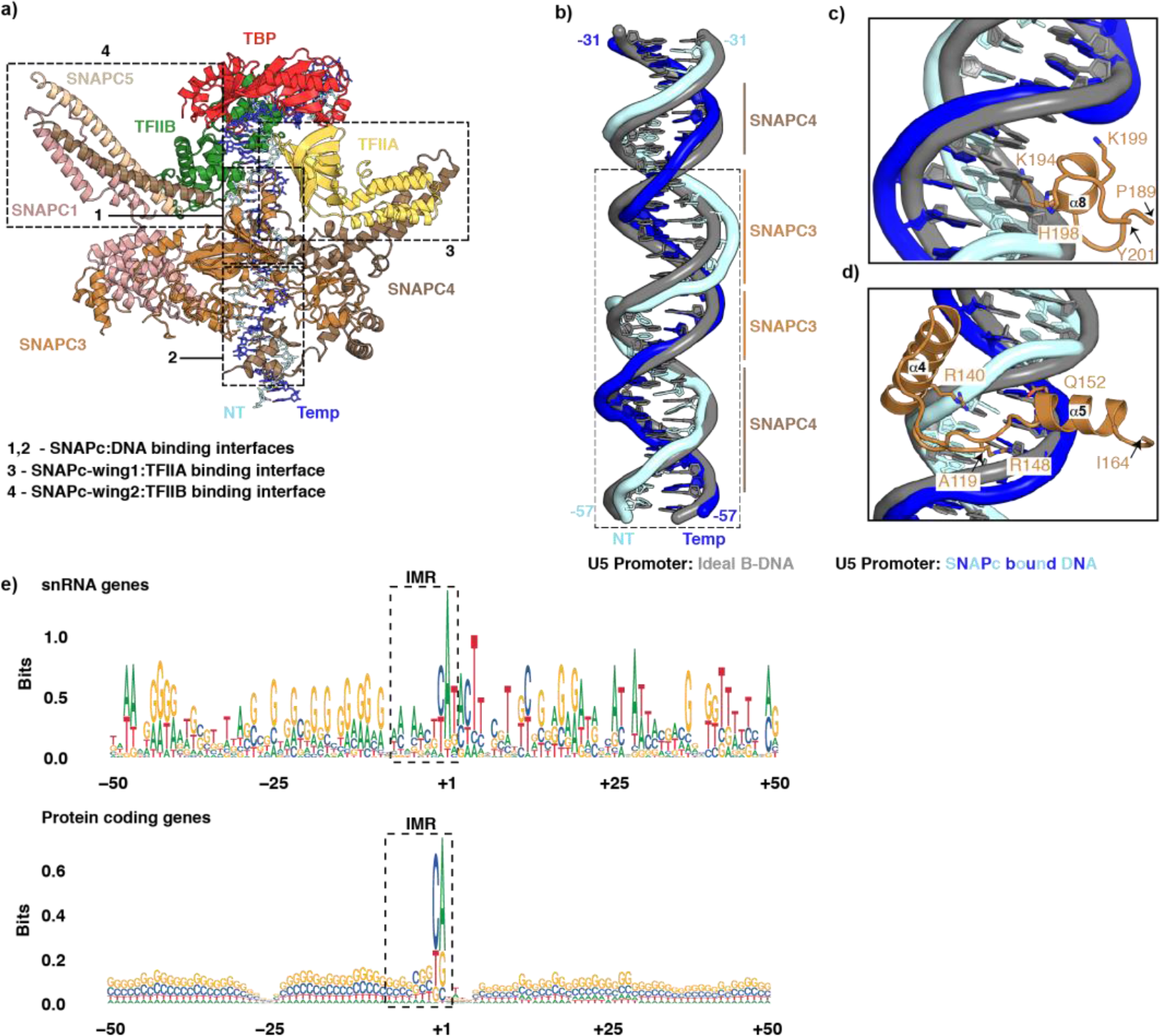
Related to Figures 4, 5 and 6. a) Birds-eye view of the SNAPc interaction with the GTFs’ and the PSE motif on U5 snRNA promoter. The dashed boxes indicate the observed interaction surfaces within the complex (1-4). b) Structural super-position of ideal B-DNA of U5 promoter to the SNAPc bound experimental DNA structure. Major and minor grooves of U5 promoter bound by SNAPC3 and SNAPC4are labelled and highlighted with lines. Dashed box indicates the PSE region. c) Close up view of SNAPC3 helix α8 binding to major groove of U5 promoter. The observed steric clash of K194 with B-DNA highlights the distortion upon SNAPc binding. d) Close up view of SNAPC3 helices α4, α5 region binding to minor groove of U5 promoter. The views in panels c and d correspond to Figure 4b, c e) Sequence logos of DNA sequence surrounding TSS peaks in expressed constitutive first/single exons for all snRNA genes (n=18) and protein coding genes (n=4721), sorted by TSS precision scores. The boxes indicate the IMR region (−8 to −2) of promoter flanking the TSS (+1). While the protein coding genes do not show any enrichment of specific nucleotides, snRNA genes present a AT-rich profile in the IMR region, indicating its tendency for spontaneous promoter opening.

## EXTENDED DATA TABLE

**Extended Data Table 1.** Cryo-EM data collection, refinement and validation statistics.

|  | RNU1-OC<br>(EMDB-xxxx)<br>(PDB xxxx) | RNU1-OC<br>Local map<br>(EMDB-xxxx) | RNU1-CC<br>(EMDB-xxxx)<br>(PDB xxxx) | RNU5-CC<br>(EMDB-xxxx)<br>(PDB xxxx) | RNU5-CC<br>Local map<br>(EMDB-xxxx) |
| --- | --- | --- | --- | --- | --- |
| **Data collection and processing** |  |  |  |  |  |
| Magnification |  | 81,000x |  | 81,000x |  |
| Voltage (kV) |  | 300 |  | 300 |  |
| Electron exposure (e-/Å<sup>2</sup>) |  | 54.45 |  | 51.93 |  |
| Defocus range (μm) |  | -0.5 to -3.0 |  | -0.5 to -2.5 |  |
| Pixel size (Å) |  | 1.05 |  | 1.05 |  |
| Micrographs collected |  | 16,854 |  | 4,842 |  |
| Initial particle images (no.) |  | 5,181,947 |  | 1,299,523 |  |
| Final particle images (no.) | 137,246 | 137,246 | 47,293 | 85,787 | 85,787 |
| Map resolution (Å) | 3.0 | 3.5 | 3.4 | 3.0 | 3.2 |
| FSC threshold | 0.143 | 0.143 | 0.143 | 0.143 | 0.143 |
| Map resolution range (Å) | 2.8 – 5.2 | 3.4 – 6.0 | 3.0 – 7.0 | 2.75 – 5.25 | 3.0 – 5.0 |
| **Refinement** |  |  |  |  |  |
| Initial model used (PDB code) | 7NVU | 7NVU | 7NVS | 7NVS | 7NVS |
| Map sharpening *B* factor (Å<sup>2</sup>) | -10 | -10 | -5 | -10 | -10 |
| **Model composition** |  |  |  |  |  |
| DNA | 126 | 93 | 132 | 132 | 83 |
| Protein residues | 5842 | 1500 | 5789 | 5789 | 1500 |
| Ligands | 11 | 2 | 12 | 12 | 2 |
| ***B* factors (Å<sup>2</sup>)** |  |  |  |  |  |
| DNA | 227.04 | 173.83 | 318.34 | 248.19 | 95.70 |
| Protein residues | 111.18 | 185.97 | 215.69 | 124.47 | 109.50 |
| Ligands | 164.08 | 158.45 | 237.49 | 194.42 | 82.54 |
| **R.m.s. deviations** |  |  |  |  |  |
| Bond lengths (Å) | 0.005 | 0.003 | 0.004 | 0.007 | 0.003 |
| Bond angles (°) | 0.756 | 0.606 | 0.509 | 0.658 | 0.524 |
| **Validation** |  |  |  |  |  |
| MolProbity score | 1.77 | 1.72 | 1.69 | 1.63 | 1.67 |
| Clashscore | 9.25 | 10.84 | 9.85 | 7.55 | 7.63 |
| Poor rotamers (%) | 0.16 | 0.00 | 0.00 | 0.00 | 0.00 |
| CaBLAM outliers | 1.93 | 1.73 | 1.63 | 1.91 | 2.07 |
| Cβ outliers | 0.00 | 0.00 | 0.00 | 0.00 | 0.00 |
| **Ramachandran plot** |  |  |  |  |  |
| Favored (%) | 95.91 | 97.01 | 97.03 | 96.68 | 96.27 |
| Allowed (%) | 4.09 | 2.99 | 2.97 | 3.32 | 3.73 |
| Disallowed (%) | 0.00 | 0.00 | 0.00 | 0.00 | 0.00 |

## Notes

### Competing Interest Statement

The authors have declared no competing interest.

